# Mechanistic computational modeling of monospecific and bispecific antibodies targeting interleukin-6/8 receptors

**DOI:** 10.1101/2023.12.18.570445

**Authors:** Christina MP Ray, Huilin Yang, Jamie B Spangler, Feilim Mac Gabhann

## Abstract

The spread of cancer from organ to organ (metastasis) is responsible for the vast majority of cancer deaths; however, most current anti-cancer drugs are designed to arrest or reverse tumor growth without directly addressing disease spread. It was recently discovered that tumor cell-secreted interleukin-6 (IL-6) and interleukin-8 (IL-8) synergize to enhance cancer metastasis in a cell-density dependent manner, and blockade of the IL-6 and IL-8 receptors (IL-6R and IL-8R) with a novel bispecific antibody, BS1, significantly reduced metastatic burden in multiple preclinical mouse models of cancer. Bispecific antibodies (BsAbs), which combine two different antigen-binding sites into one molecule, are a promising modality for drug development due to their enhanced avidity and dual targeting effects. However, while BsAbs have tremendous therapeutic potential, elucidating the mechanisms underlying their binding and inhibition will be critical for maximizing the efficacy of new BsAb treatments. Here, we describe a quantitative, computational model of the BS1 BsAb, exhibiting how modeling multivalent binding provides key insights into antibody affinity and avidity effects and can guide therapeutic design. We present detailed simulations of the monovalent and bivalent binding interactions between different antibody constructs and the IL-6 and IL-8 receptors to establish how antibody properties and system conditions impact the formation of binary (antibody-receptor) and ternary (receptor-antibody-receptor) complexes. Model results demonstrate how the balance of these complex types drives receptor inhibition, providing important and generalizable predictions for effective therapeutic design.

## INTRODUCTION

Interleukin-6 (IL-6) and interleukin-8 (IL-8) play key roles in inflammation and have been implicated in cancer progression. IL-6 is a pro-inflammatory cytokine that, along with TNFα and IL-1β, contributes to the early response to infection [1]. IL-6 is produced by macrophages, dendritic cells, and epithelial cells in response to pathogens, and this cytokine drives T cell differentiation as well as plasma B cell differentiation [2]. IL-8 (CXCL8) is a chemokine produced by monocytes, macrophages, fibroblasts, and other cells [3–5], and it is responsible for attracting leukocytes (typically but not exclusively neutrophils) to the site of inflammation by enhancing extravasation and by chemoattraction within tissue [4].

Both IL-6 and IL-8 have been implicated in the pathogenesis and progression of several solid tumor types, including breast [6], prostate [7], colon [8], and pancreatic [9] cancers, and indeed elevated expression of both molecules has been associated with increased cancer aggressiveness and metastatic burden [8,10,11]. It has recently been demonstrated that IL-6 and IL-8 paracrine signaling in metastatic cancer cells increases motility in a cell density-dependent manner [12]. Above a threshold cell density, cancer cells secrete IL-6 and IL-8, which synergistically activate a complex paracrine signaling pathway via Janus kinase (JAK2) and signal transducer and activator of transcription 3 (STAT3) that prompts cancer cells to form *Arp2/3*-dependent dendritic protrusions and undergo migration. Simultaneous inhibition of the IL-6/IL-8 signaling network with tocilizumab, a monoclonal anti-IL-6 receptor-alpha (IL-6Rα, hereafter denoted IL-6R) antibody primarily used to treat rheumatoid arthritis [13], and reparixin, a small molecule allosteric inhibitor of the IL-8 receptor (IL-8R) that recently completed phase II clinical trials against breast cancer [14], was found to decrease *in vitro* cell migration and significantly decrease *in vivo* metastasis without affecting rates of tumor growth [12,15]. Collectively, these data suggest that targeting IL-6, IL-8, and their receptors is a promising approach to inhibiting tumor metastasis and cancer lethality.

Monoclonal antibodies (mAbs) have been a fixture in anti-cancer therapeutic regimens over the past 20 years [16,17] due to their target specificity, *in vivo* stability, modular construction, and multi-faceted actions. However, monospecific mAbs have limitations, including the emergence of acquired resistance as cancer cells mutate [18]. Bispecific antibodies (BsAbs), antibodies engineered to simultaneously engage two different target molecules, demonstrate great potential to overcome the shortcomings of antibody drugs [19–21]. Binding in *cis*, i.e., with the BsAb bridging different receptors on the same cell, confers avidity, improved tissue selectivity, and reduced off-target side effects [22–24], while also reducing the likelihood of drug resistance [25]. Additionally, concurrent binding to separate targets can prevent receptor homodimerization [26] and increase treatment potency in target tissues [27].

To study and potentially better target the IL-6/IL-8 signaling network for metastasis inhibition, Yang and colleagues recently engineered a novel bispecific antibody, BS1, against IL-6R and IL-8RB (also known as CXCR2, hereafter called IL-8R) [28]. BS1 contains arms with variable domains of two distinct antibodies: the anti-IL-6R antibody tocilizumab and the anti-IL- 8R antibody 10H2 [5,29] [**Figure 1A**]. BS1 significantly reduced *in vitro* cancer cell migration, effecting greater inhibition of migration than either the combination of tocilizumab and reparixin or the combination of tocilizumab and 10H2 [28]. Furthermore, BS1 potently decreased metastatic burden *in vivo* in orthotopic mouse xenograft models and, when paired with the anti-proliferative agent gemcitabine, significantly decreased both metastasis and tumor growth [28]. In all studies, BS1 outperformed combination treatments, demonstrating the effectiveness of bispecific agents in targeting complex signaling networks.

**Figure 1.**
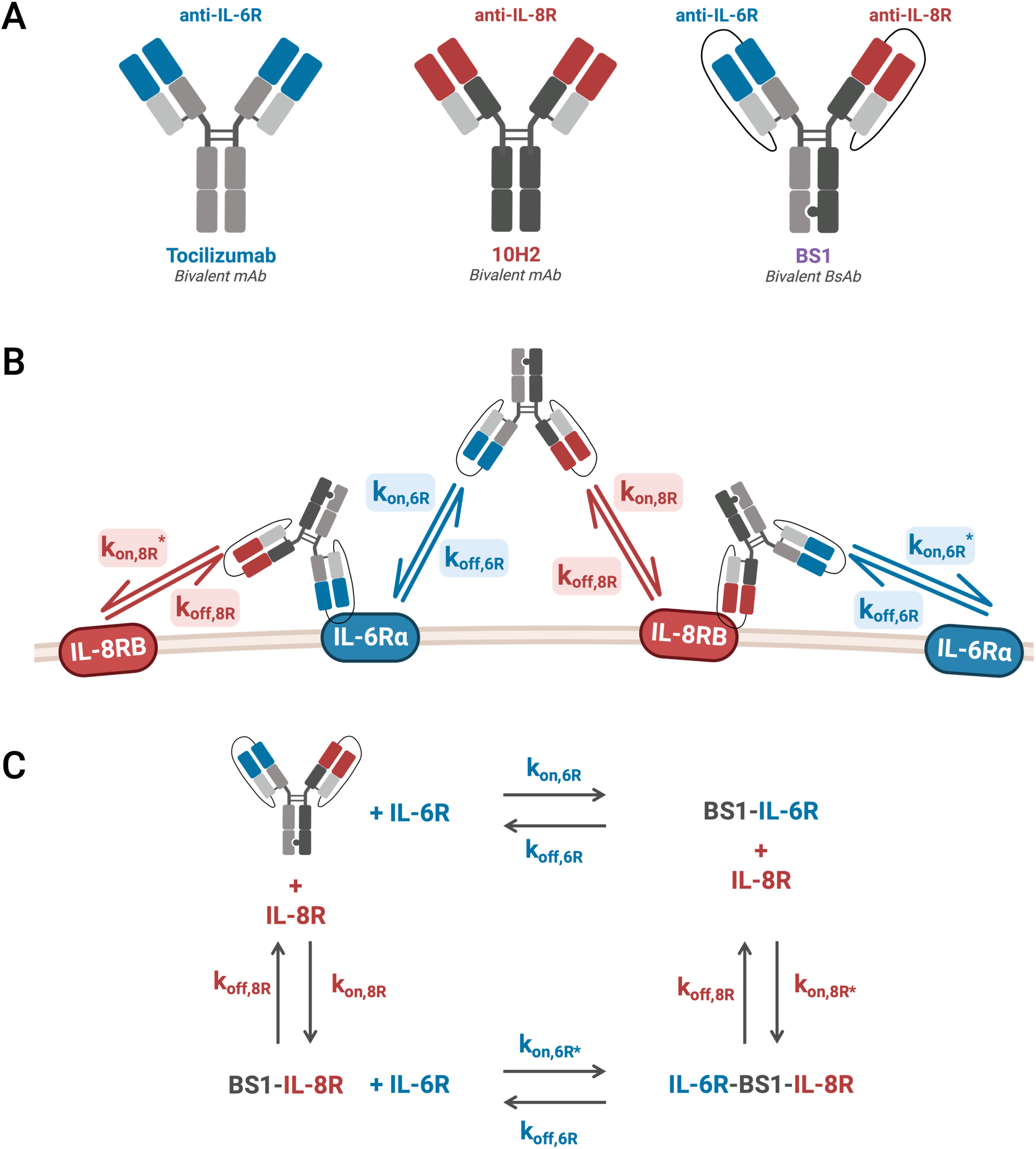
Bivalent antibody binding model antibodies, rate constants, and reactions. **A,** Monoclonal (mAb) and bispecific (BsAb) antibodies simulated in our computational model. Tocilizumab is a recombinant humanized mAb with two anti-IL-6Rα (denoted anti-IL-6R) binding domains; 10H2 is a mAb with two anti-IL- 8RB (denoted anti-IL-8R) binding domains; BS1 is an anti-IL-6Rα/anti-IL-8RB BsAb synthesized from the binding domains of tocilizumab and 10H2 by combining the knobs-into-holes and single-chain Fab methodologies. **B,** Schematic of the IL-6Rα/IL-8RB/BS1 antibody-binding model kinetics. BS1 can bind to either IL-6Rα or IL-8RB, and, having done so, the BS1-receptor complex can then bind to the other receptor. kon,6R and kon,8R describe the association rates for the formation of binary antibody-receptor complexes, and kon,6R* and kon,8R* describe the association rates for the formation ternary receptor-antibody-receptor complexes. The same koff,6R and koff,8R rate constants are used for the dissociation of both the binary and the ternary complexes. Schematics for the two monoclonal antibodies, tocilizumab and 10H2, are included in the *Supplemental Information* [**Figure S1**]. **C,** Simplified view of the schematic in **B** illustrates how the reactions form a thermodynamic cycle. The reactions can proceed in a clockwise or counter-clockwise manner to return back to the starting reactants, forming a cycle with a net free energy change of 0. This figure was created with BioRender.com.

However, while the dual-targeting ability of BsAbs holds promise, it will be crucial to understand the mechanisms underpinning BsAb binding and how those mechanisms differ from treatment with a combination of monospecific mAbs in order to maximize efficacy. As antibodies are multivalent, their binding is driven both by the inherent affinity of each binding domain for its target antigen and by avidity, the accumulated binding strength from each of the individual molecular interactions [30–32]. While it has been established that avidity plays a key role in BsAb tissue selectivity and therapeutic efficacy [23,33], the interplay of individual domain affinity, overall avidity, target expression, and therapeutic concentration in the context of cell binding remains poorly understood. Mechanistic computational models of antibody-target interactions can address this knowledge gap—by incorporating parameters for both monovalent binding affinity and multivalent binding avidity with differential equations describing the binding kinetics, we can investigate the influence of these factors on the binding of monospecific and bispecific antibodies [34–36].

Mathematical models of the kinetics of heterobivalent antibody binding to cell surface targets have been characterized previously [24,37–41], providing a general framework for modeling multivalent binding. Here, we extend existing modeling frameworks to build a quantitative, computational model for a specific therapeutic target: antibodies targeting the IL-6 and IL-8 receptors for the prevention of cancer metastasis. To therapeutically inhibit IL-6/IL-8- mediated metastasis, we are examining two existing monospecific antibodies, tocilizumab (denoted anti-IL-6R) [42] and 10H2 (denoted anti-IL-8R) [5,29], and the novel bispecific antibody, BS1 (denoted anti-IL-6R/anti-IL-8R), first described by Yang et al. [28] [**Figure 1A**]. Binding mechanics (receptor complex formation) of each antibody are expressed in a series of coupled differential equations, creating a complete model of antibody interactions that is faithful to the biophysics.

Our detailed simulations of the *in vitro* monovalent and bivalent binding interactions between different antibody constructs and the target receptors, IL-6R and IL-8R, establish how binary (antibody-receptor) and ternary (receptor-antibody-receptor) complex formation drives target inhibition. We demonstrate that the ratio between expression levels of IL-6R and IL-8R is crucial to bispecific antibody binding *in vitro* and leads to significant differences in monospecific and bispecific antibody behavior. Simulations also predict necessary antibody concentrations for optimal binding, namely that the most stable antibody-receptor complex formation occurs at high receptor concentrations and intermediate antibody concentrations. Overall, our model simulations with different antibody constructs clarify the effects of binding domain affinity and target expression on receptor inhibition, providing insight that is applicable not only to our particular BsAb system, but also more broadly to bispecific antibody therapeutic design.

## METHODS

### Monospecific and Bispecific Antibodies

The three key antibodies of interest we are using to target the receptors in the IL-6/IL-8 system are tocilizumab (anti-IL-6Rα), 10H2 (anti-IL-8RB), and a novel bispecific antibody developed by Yang et al. [28], BS1 (anti-IL-6Rα/anti-IL-8RB) [**Figure 1A**]. BS1 is a human immunoglobulin G (IgG)-based bispecific antibody synthesized by combining the knobs-into-holes strategy [43] with single-chain Fab design [44], and it was developed to increase high-affinity selective targeting of IL-6R and IL-8R, decrease off-target toxicity, and reduce risk of acquired resistance [28]. The IL-6Rα-blocking arm of BS1 comes from tocilizumab, a monoclonal anti-IL-6Rα antibody used to treat rheumatoid arthritis [13]. There are no clinically-approved anti-IL-8R antibodies, but the experimental anti-IL-8RB antibody 10H2 blocks IL-8 binding and activity [5,29,45] and is used for the IL-8RB-blocking arm of BS1. BS1 is bivalent (one anti-IL-6Rα domain and one anti-IL- 8RB domain) and interacts with IL-6Rα^+^/IL-8RB^+^-transduced HEK 293T cells in flow cytometry-based binding studies (KD = 14.4 nM) [28].

### Binding Model Equations

To describe the ligand-receptor and antibody-receptor binding kinetics, we built a coupled set of ordinary differential equations (ODEs) using the law of mass action. Each individual ODE describes one molecule or molecular complex, with terms representing each binding interaction (binding and unbinding processes) in the system [**Figures 1B, S1**].

The equations take the form:

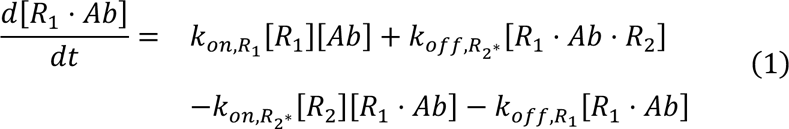

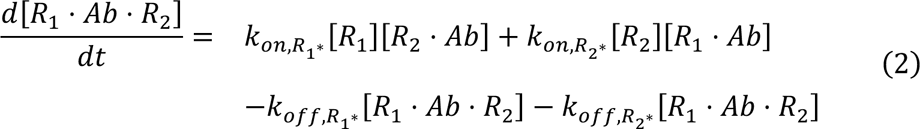

These equations represent the antibody first binding one receptor and then binding a second receptor (of the same or different type, depending on the antibody). The “second binding” events, describing the cross-linking of an antibody-receptor complex with an additional receptor, are indicated by asterisks in the equations. The full model equations are included in the *Supplemental Information*. The experimental data to which we are comparing our model simulations [28] were acquired at 4°C. As a result, other processes that could have been incorporated into the model, including receptor synthesis, internalization, and degradation, were assumed to be negligible because they are typically suppressed at low temperatures.

### Rate Constant Values and Similarity of Binding Sites

To simulate the complete mechanistic ODE model (see *Supplemental Information*) for three different antibodies requires values for the many parameters in the model—primarily, rate constants. The number of unique parameters for which we need values can be reduced using (a) the identification of a thermodynamic cycle and (b) assumptions of the similarity of binding sites across antibodies, as described in the next two sections.

### BsAb-Receptor Binding Reaction Cycle

The bivalent antibody reactions function in a cycle where all of the species are linked by the reactions. Given the reactions:

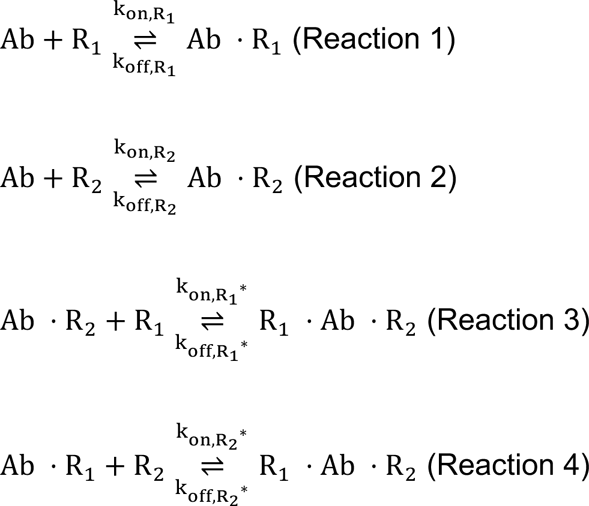

These reactions form a cycle because it is possible to find a path through the reactions that leads back to the starting point. For example, starting with the antibody in Reaction 1, the forward reaction (binding) produces Ab⋅R1. Then, the forward reaction in Reaction 4 creates R1⋅Ab⋅R2. Using the reverse reaction in Reaction 3, Ab⋅R2 is obtained. Finally, through the reverse reaction in Reaction 2, the original antibody is returned. Because there is a path through the reactions that returns the original reactant and does not repeat any reactions, the reactions form a cycle, with the forward reactions of Reactions 1 and 4 and the reverse reactions of Reactions 2 and 3.

For each of these reactions, at equilibrium, the rate of the forward reaction must be equal to the rate of the reverse reaction by the principle of detailed balance [46,47]. Thus, for each reaction, the concentrations at equilibrium and the rate constants can be related:

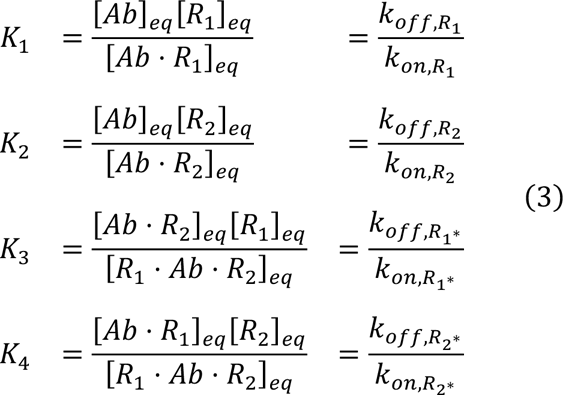

To relate the rate constants for the full reaction cycle, we can multiply the equilibrium constants for each reaction, using the inverse equilibrium constant for each reaction that is reversed (Reactions 2 and 3):

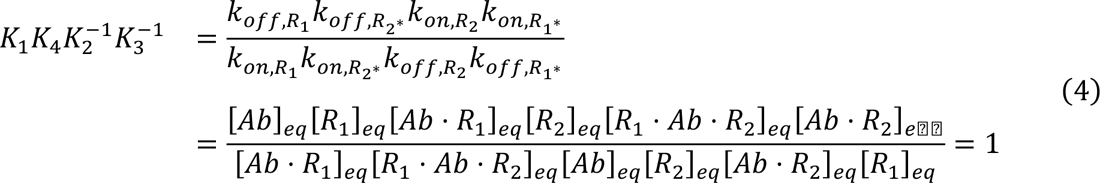

All of the concentrations in Equation 4 cancel out, and the product of the equilibrium constants for the cycle reactions is unity. This is an established behavior for cycles where ligand is bound and released into a single volume with no other reactions [48,49]. Although Equation 4 was derived from the equilibrium relationships, the result only involves the system constants, and thus it applies even when the system is not in equilibrium [48].

### Similarity of Binding Sites

Using the relationship of the equilibrium constants to the rate constants, we can rewrite the cycle constraint:

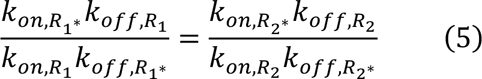

Due to the similarity of the dissociation reaction kinetics between the different antibody domains [50], the parameter space can be simplified by assuming that the koff values for the first binding and second binding reactions are equal (i.e., koff,R1 = koff,R1* and koff,R2 = koff,R2*). This assumption further simplifies the rate constant relationship to:

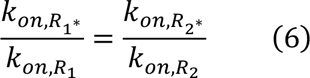

In their bivalent antibody model, Harms et al. termed this ratio of kon,R* to kon,R as the “cross-arm binding efficiency” (χ) [38,51]. This value combines multiple factors that impact multivalent binding, including the increased rate of binding due to the restriction of the bound antibody to a small volume adjacent to the cell membrane and the decreased flexibility and rotational freedom of the tethered antibody. Greater values of χ indicate stronger cross-arm binding, leading to greater rates of ternary complex formation.

With the reaction cycle constraint and above koff assumption, the number of unique parameters needed to describe an individual bivalent antibody is reduced from eight to five— three kon values and two koff values.

We can further simplify the overall number of parameters describing the binding of the three antibodies being studied here by noting that BS1 is synthesized using the IL-6Rα and IL- 8RB binding domains of tocilizumab and 10H2, respectively. Thus, we made an additional simplifying assumption that the association rate constants were equal for the similar binding domains. With this assumption, the independent association rate constants become:

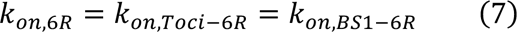

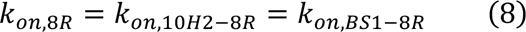

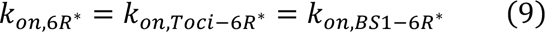

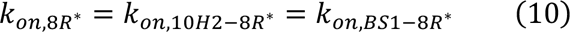

With the previous parameter space reduction from the binding reaction cycle constraint and the assumption that first and second koff values are equal, the full parameter space for tocilizumab, 10H2, and BS1 is reduced to five independent parameters: kon,6R, kon,8R, kon,6R*, koff,6R, and koff,8R, with kon,8R* being dependent on the values of the other association rates.

### IL-6Rα and IL-8RB HEK 293T Cell Surface Binding Assays

The association and dissociation rate constants for the antibody-receptor complexes were estimated here using experimental data from *in vitro* cell surface binding flow cytometry assays; these data were previously reported [28] and detailed methods can be found in the original publication. Briefly, the IL-6Rα and IL-8RB genes were transduced into HEK 293T cells, generating four cell lines: IL-6Rα^+^/IL-8RB^-^, IL-6Rα^-^/IL-8RB^+^, IL-6Rα^+^/IL-8RB^+^, and IL-6Rα^-^/IL- 8RB^-^. The receptor expression on the transduced cell lines was quantified through flow cytometry [**Table 1**].

**Table 1.** IL-6Rα and IL-8RB quantification on transduced HEK 293T cells (expressed in # receptors/cell).

| Cell Line | IL-6R Expression | IL-8R Expression |
| --- | --- | --- |
| IL-6Rα<sup>+</sup>/IL-8RB<sup>-</sup> | $5.08 \times 10^5$ | N/A |
| IL-6Rα<sup>-</sup>/IL-8RB<sup>+</sup> | N/A | $1.30 \times 10^6$ |
| IL-6Rα<sup>+</sup>/IL-8RB<sup>+</sup> | $3.16 \times 10^5$ | $6.18 \times 10^5$ |

The transduced cell lines were placed into 96-well plates (1 × 10^5^ cells per well) and incubated with doses of monoclonal or bispecific antibodies (tocilizumab, 10H2, and BS1) from 10^-2^ to 10^3^ nM for two hours at 4°C. Cells were then washed and incubated with an allophycocyanin (APC)-conjugated anti-human IgG1 antibody for 15 min at 4°C. Antibodies binding to the receptors were quantified with flow cytometry and reported as Mean Fluorescent Intensity (MFI) detected. The experiments were performed with three technical repeats, and data from all three replicates was used for the binding parameter optimization.

### Model Parameterization: Optimization of Binding Rate Constants

We fit the antibody-receptor binding model to the flow cytometry binding assay data by estimating the association and dissociation rate constant values for each binding step. To reduce the number of parameters required to fully characterize the model, multiple simplifying assumptions were made about similarities in binding domain structure and protein geometry, and from the thermodynamic cycle constraint, as described in the previous sections. Thus, to describe the system fully for tocilizumab, 10H2, and BS1, we need values for five unique parameters: kon,6R, kon,8R, kon,6R*, koff,6R, and koff,8R, with the value of kon,8R* being dependent on the values of the other parameters [**Figures 1B, 1C**].

We created MATLAB code to describe the ordinary differential equation (ODE) model as a system of equations and used the ode15s solver to simulate the system over time. Simulations were performed under conditions replicating the *in vitro* experiments as closely as possible. For example, the initial concentration of receptors for each cell line was set to values from the transduced HEK293T cells [**Table 1**], and the initial antibody concentrations used in the simulation were the same as the range of the experimental values. The antibody was added at time = 0, and the free antibody concentration was set to 0 at time = 2 hours to simulate the washing out of unbound antibody. The simulation was continued for 15 minutes after the washout to mimic the incubation with the APC-conjugated antibody. For the optimization, the total bound antibody at the final simulation time point was compared to the Mean Fluorescent Intensity (MFI) values from the flow cytometry binding assays, which represent the total bound antibody in that experiment.

We used the non-linear least squares optimization function lsqnonlin to determine the parameter values (i.e., rate constant values) that minimized the sum of the squared differences between the simulation output and the experimental data points. Three hundred sets of initial guesses for the parameter values were generated using Latin Hypercube Sampling, using a log-uniform distribution for each parameter over the ranges listed in **Table 2**. The optimization process was repeated for each set and for each type of simulation normalization (described below), and optimizations that did not converge or that did not vary from the initial guesses were discarded (approximately 13% of the total optimization runs).

**Table 2.** Binding rate constants estimated from IL-6Rα and IL-8RB HEK 293T cell surface binding assays. S.D. = standard deviation of log-transformed optimized values. α = 8.3 × 10^-7^ nM/(# / cell) is used to convert the rate constants between nM and number of receptors per cell. The derivation of this value for the unit conversion is included in the Supplemental Information.

| Parameter | Initial Guess | Optimization |  | S.D. | Units | Reference |
| --- | --- | --- | --- | --- | --- | --- |
|  | Range | Bounds | Best Fit Value |  |  |  |
| $k_{on,6R}$ | $[10^{-9}, 10^{-2}]$ | $[10^{-11}, 1]$ | $5.92 \times 10^{-6}$ | 1.15 | $nM^{-1}s^{-1}$ | Best fit to the data |
| $k_{on,8R}$ | $[10^{-9}, 10^{-2}]$ | $[10^{-11}, 1]$ | $9.03 \times 10^{-6}$ | 0.953 | $nM^{-1}s^{-1}$ | Best fit to the data |
| $k_{on,6R^*}$ | $[10^{-13}, 10^{-5}]$ | $[10^{-15}, 10^{-3}]$ | $8.11 \times 10^{-8}$ | 2.10 | $\left(\frac{\#}{cell}\right)^{-1} s^{-1}$ | Best fit to the data |
| $\alpha^{-1} \cdot k_{on,6R^*}$ | $[1.2 \times 10^{-7}, 12]$ | $[1.2 \times 10^{-9}, 1.2 \times 10^3]$ | $9.77 \times 10^{-2}$ | 2.10 | $nM^{-1}s^{-1}$ | $k_{on,6R^*}$ converted to $nM^{-1}s^{-1}$ |
| $k_{on,8R^*}$ | — | — | $1.24 \times 10^{-7}$ | 2.25 | $\left(\frac{\#}{cell}\right)^{-1} s^{-1}$ | Dependent on other binding constants |
| $\alpha^{-1} \cdot k_{on,8R^*}$ | — | — | $1.49 \times 10^{-1}$ | 2.25 | $nM^{-1}s^{-1}$ | $k_{on,8R^*}$ converted to $nM^{-1}s^{-1}$ |
| $k_{off,6R}$ | $[10^{-7}, 10^{-2}]$ | $[10^{-9}, 1]$ | $5.61 \times 10^{-5}$ | 1.08 | $s^{-1}$ | Best fit to the data |
| $k_{off,8R}$ | $[10^{-7}, 10^{-2}]$ | $[10^{-9}, 1]$ | $6.38 \times 10^{-5}$ | 1.36 | $s^{-1}$ | Best fit to the data |

The MFI values from the flow cytometry binding assays were normalized against the values for bound BS1 at the initial antibody concentrations where binding reached saturation. The MFI values for BS1 where binding reached saturation were averaged and used as the denominator to normalize all of the data for all antibodies in a single cell line; each cell line was normalized separately. For the simulation results, four different normalization schemes were tested in the parameter optimization; these normalizations are summarized in **Table 3** and are defined as follows. *BS1* indicates simulations that were normalized against the amount of bound BS1 in the corresponding cell line, and *Ab* indicates simulations where each antibody was normalized against the amount of that specific antibody bound in the corresponding cell line. *Data* indicates simulations that were normalized using output at the same concentrations that were used to normalize the experimental data, and *Max* indicates simulations that were normalized using the output at the maximum antibody concentration.

**Table 3.** Normalization schemes tested in the optimization of the binding rate constants against the flow cytometry binding assay data. The “BS1, Data” scheme was selected for the parameter optimization.

| **Scheme** | **Normalized Against** | **Initial Concentrations** |
| --- | --- | --- |
| BS1, Data | BS1 | Concentrations used for experimental data |
| BS1, Max | BS1 | Maximum antibody concentration |
| Ab, Data | Individual Ab | Concentrations used for experimental data |
| Ab, Max | Individual Ab | Maximum antibody concentration |

As discussed in the *Results*, the optimizations performed with normalization against BS1 at binding saturation (labeled “BS1, Data”) show stronger convergence around a single optimal value for each parameter and less dependence on the initial value used, so this normalization scheme was selected for the parameter optimization. The cost of the parameter set was calculated as the sum of the squared difference between the normalized model output and normalized experimental binding at each input concentration. The best fit parameter set was used for further binding model simulations [**Table 2**].

### Simulation Output

As described above, for the optimization of the binding parameters, the total receptor-bound antibody at the final simulation time point was output from the model simulations for comparison to the experimental MFI values, which represent the total bound antibody in the flow cytometry assays. For the subsequent model simulations, as bivalent antibodies can form both binary (antibody-receptor) and ternary (receptor-antibody-receptor) complexes, quantifying the amount of antibody-bound receptor (as distinct from receptor-bound antibody) provides more information about the inhibition of the system. Thus, for the simulations using the parameterized model, we output the concentration of antibody-bound receptor (in # receptors/cell). Results are given as the concentration of receptor bound in a particular complex type, either binary complexes (with a single receptor) [**Equation 11**] or ternary complexes (with two receptors) [**Equation 12**], or the total bound receptor [**Equation 13**], which is the sum of receptor bound in binary and ternary complexes. Results are also shown for the receptor fractional occupancy, which is calculated as the fraction of the total receptor concentration in the system that is bound either in a specific complex type or overall. Unless stated otherwise, the bound receptor refers to the sum of IL-6R and IL-8R bound in a particular complex type or overall.

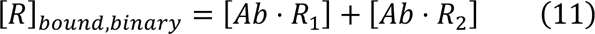

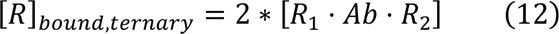

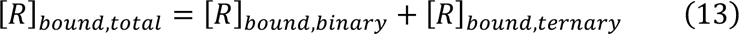

### Univariate Sensitivity Analysis

We performed local and global univariate sensitivity analyses of the binding model rate constants and the species concentrations to determine the effects of the individual parameters on the model output. For the local sensitivity analysis, simulations were performed for two hours after antibody dosing, with a baseline BS1 concentration of 10 nM and baseline receptor concentrations of 5 × 10^4^ receptors/cell for both IL-6R and IL-8R. Each rate constant and initial concentration was varied 10 percent above its baseline value, with each parameter varied individually in separate simulations. The Area Under the Curve (AUC), calculated as the integration of the BS1-receptor complex concentration over time, was output for both ternary complexes and total bound receptors. The sensitivity for each parameter-output combination was calculated as the percentage change in the output value divided by the percentage change in the parameter value (10 percent for all simulations).

In the global sensitivity analysis, simulations were performed for 24 hours after antibody dosing, with constant receptor concentrations of 5 × 10^4^ receptors/cell for both IL-6R and IL-8R. The longer simulation time was selected for this analysis to examine the model output closer to equilibrium. Each association and dissociation rate constant was separately varied over two orders of magnitude below and above its optimized value. The receptor fractional occupancy was output for ternary complexes and total bound receptor, with fractional occupancy calculated as the fraction of the total receptor (IL-6R + IL-8R) that is bound in ternary complexes or bound in total in either binary or ternary complexes.

## RESULTS

### Binding Parameter Optimization and Parameter Identifiability

The optimization of the association and dissociation rate constants for the antibody-receptor binding model generated a range of optimal parameter sets depending on the initial guesses used [**Figure 2A**]. About half of the optimized parameter sets, and in particular those of the lowest cost (i.e., best fit), resulted in consistent parameter values (horizontal patterns on the graph) that are independent of the initial guess values. Some of the optimization results do give parameter sets that are correlated to initial guesses, but these are higher in cost (i.e., poorer fit overall) and fewer in number; since each point is moderately transparent in the graph, darker regions indicate many overlapping optimized values. All five of the parameters optimized show consistent optimal parameter values obtained from a wide range of initial guesses. This is evidence of good parameter identifiability— given five parameters being optimized against seven sets of data points from experiments across three cell types and three antibodies.

**Figure 2.**
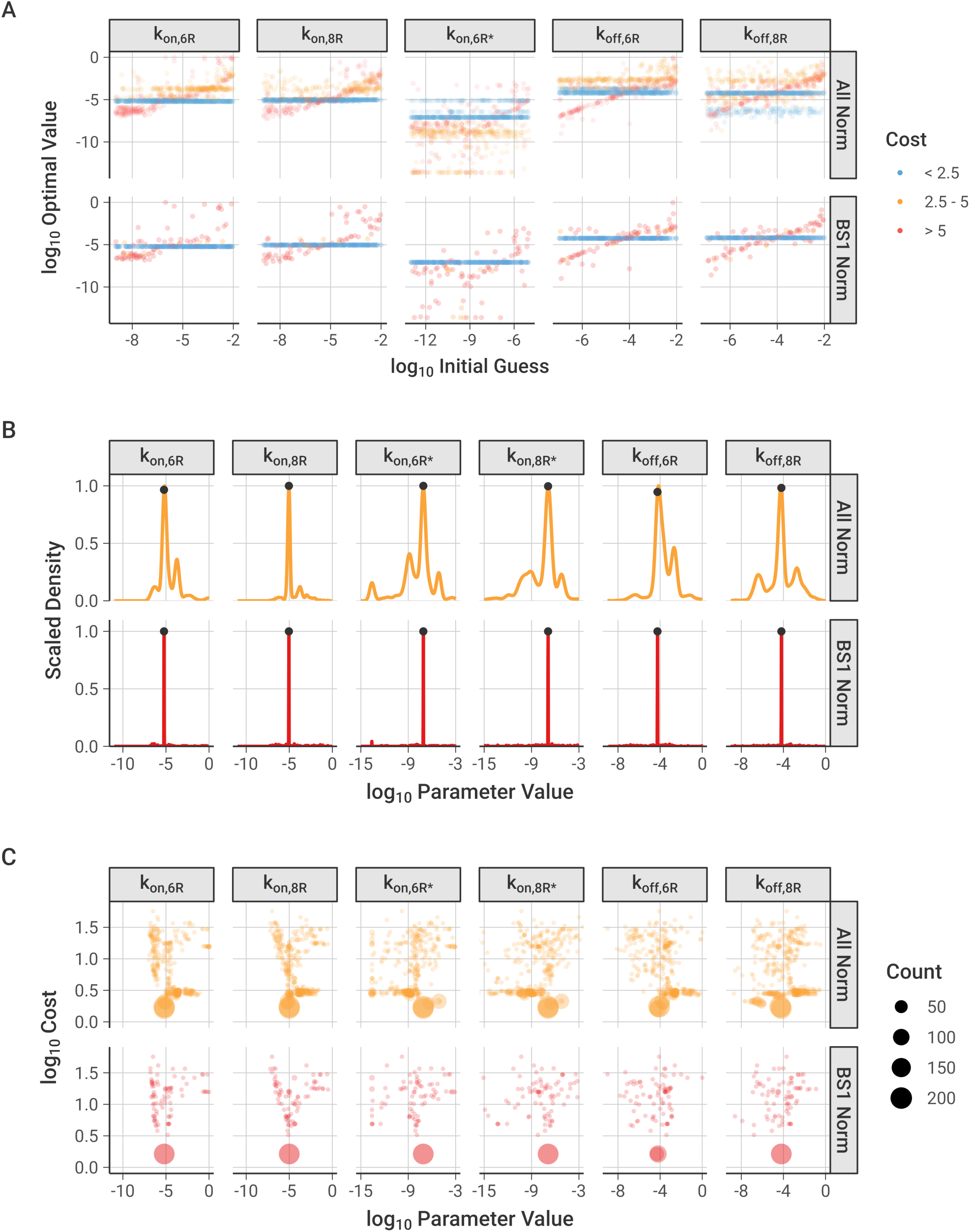
Optimization of binding association and dissociation constants to experimental data. The cost function is calculated as the sum of the squared differences between the normalized model output and the normalized experimental data at each antibody concentration used. “All Norm” includes all of the optimized parameter sets from each of the different normalization options described in the Methods, and “BS1 Norm” highlights the parameter sets where the model output was normalized against the bound concentration of BS1 at the initial concentrations used to normalize the experimental data, which was the best-performing normalization. Figures separated by normalization scheme and figures with a narrow range of parameter values are available in the *Supplemental Information* [**Figures S2-S4**]. **A,** Relationship between initial guesses and optimized values for each binding reaction rate constant. kon,8R* is not pictured because its initial and ‘optimized’ values were determined from the other parameters using the thermodynamic cycle relationship. **B,** Distribution of optimized parameter values across all optimizations performed. Marked points indicate the values of the lowest cost parameter set (values are listed in **Table 2**). **C,** Relationship between optimized parameter values and the cost of the optimized parameter sets compared to experimental data, separated by parameter. Optimized points with the same value are grouped into a single point, with the point size indicating how many optimized parameter values are in the group.

We tested whether the choice of simulation normalization scheme, as described in the *Methods*, influenced the optimization. The optimizations performed with normalization against BS1 show stronger convergence around a single optimal value for each parameter and less dependence on the initial value used [**Figure 2A, lower panel**], compared to simulations normalized to results for each antibody individually, which show a wider spread in the optimal values obtained and a greater reliance on the value of the initial guess [**Figures S2, S4**]. Further discussion of the results from the different normalization schemes is included in the *Supplemental Information*.

The frequency distributions of the optimized parameter values are narrow, again supporting good parameter identifiability and indicating that the parameters are well-constrained by the data [**Figures 2B, S3**]; the location of the best-fit (i.e., lowest cost) value is marked for each parameter. The parameter distributions contain multiple small peaks, expressing separate reoccurring optimal values, but each parameter demonstrates a distinct, most frequent value that also corresponds with the lowest cost value. Of note, the first association steps (kon) are better constrained, while the second steps (kon*) have a slightly larger range of potential values [**Table 2**].

Across the normalization schemes tested, the most frequent parameter set corresponded well with the lowest cost parameter set [**Figure 2C**]. In this visualization, optimized parameters with the same value and cost were grouped into a single point, and the area of the point was scaled with the number of parameters in the group – in other words, large dots represent parameter value-cost pairs that occur more frequently across the 300 optimizations. All six parameters (with five parameters being optimized and kon,8R* being calculated from the thermodynamic cycle constraint) show an optimal lowest-cost point that occurs most frequently, with a spread of less frequently occurring values around this central value.

Based on its low dependence on initial guess, high proportion of low-cost optimal parameter sets, and narrow distribution of optimal values, the normalization scheme with BS1 data at binding saturation was selected as the primary normalization method for the remaining analysis. The lowest cost parameter set from the optimizations performed with this normalization scheme was selected for the binding model parameter values [**Table 2**] and is indicated on the frequency distribution [**Figure 2B**].

### Best-Fit Parameters Recapitulate Experimental Observations

The best-fit association and dissociation constants [**Table 2**] fall within the range of typical values for antibody binding [52]. The association rates for binding to IL-6R and IL-8R are very close in value, and there is not a substantial difference in the monovalent binding domain affinities for either receptor [**Table 4**], as was observed experimentally [28]. Moreover, the calculated monovalent affinities are consistent with the results from the experimental characterization of BS1 [**Table S2**] [28]. The second binding step, where the binary antibody-receptor complex cross-links with an additional receptor to form a ternary receptor-antibody-receptor complex, is substantially faster than the initial binding. This is expected for bivalent binding, as the antibody is tethered to the cell surface and held in close proximity to the membrane receptors, promoting interaction with a second receptor. The cross-linking equilibrium constants are in the sub-picomolar range, and the ratio of kon,R* to kon,R, sometimes termed the “cross-arm binding efficiency” [38], is 1.6 × 10^4^, indicating strong avidity of BS1 binding.

**Table 4.** Dissociation equilibrium constants for the initial binding to form binary complexes and the cross-linking to form ternary complexes. Values are calculated from the binding rate constants [Table 2] via KD = koff / kon. The antibodies share the same equilibrium constants due to the assumptions made in the parameter optimization.

| Equilibrium |  |  |
| --- | --- | --- |
| Constant | Description | Value (nM) |
| $K_{D,6R}$ | Initial binding to IL-6R for tocilizumab and BS1 | 9.5 |
| $K_{D,8R}$ | Initial binding to IL-8R for 10H2 and BS1 | 7.1 |
| $K_{D,6R^*}$ | Cross-linking to IL-6R for tocilizumab-IL-6R and BS1-IL-8R | $5.7 \times 10^{-4}$ |
| $K_{D,8R^*}$ | Cross-linking to IL-8R for 10H2-IL-8R and BS1-IL-6R | $4.3 \times 10^{-4}$ |

Simulations using this best-fit parameter set indeed recreate the experimental *in vitro* flow cytometry data (i.e., binding of antibody to the cell surface) that was used to fit the parameter values very well [**Figure 3**]. The dose-dependence, antibody-dependence, and cell-type-dependence (i.e., receptor-expression dependence) of the data were all captured in the simulation.

**Figure 3.**
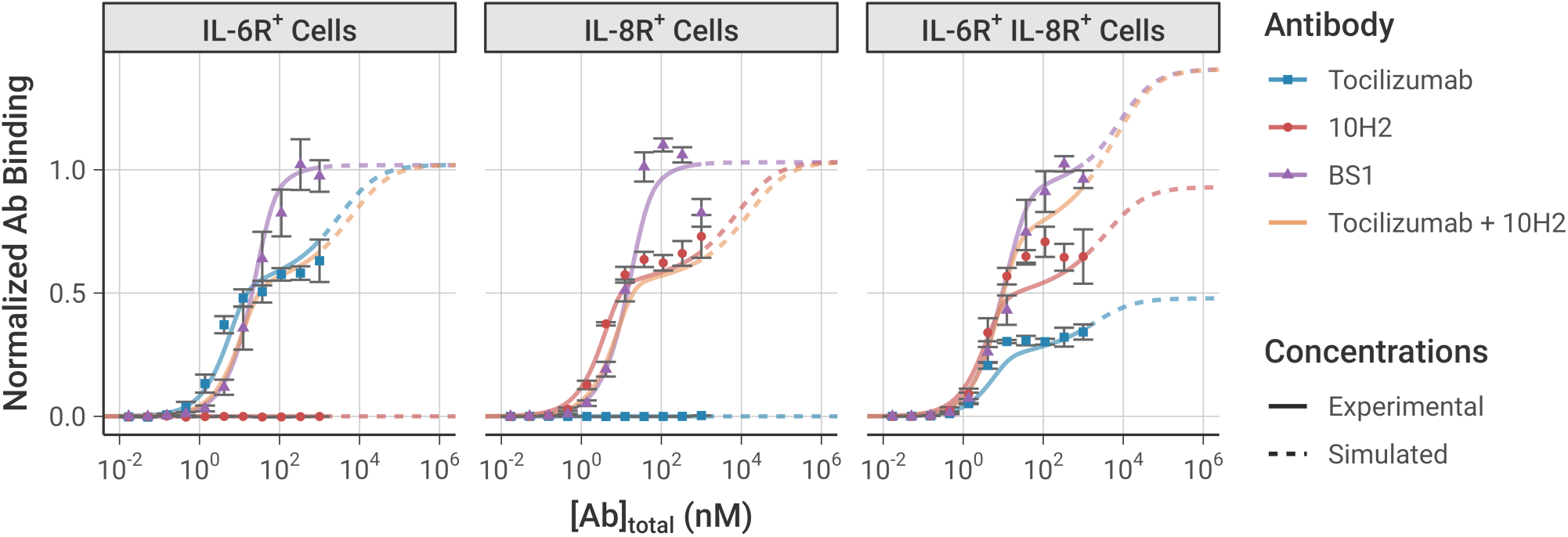
Model simulation results using the best-fit parameter set compared to the experimental data used to fit the model parameters. Simulations were performed under the same conditions as the experiment: 10^5^ cells/well, receptor expression levels from the transduced cell lines [**Table 1**], and with a 2-hour initial association period followed by a 15-minute free antibody washout. The model simulation results (lines) are compared to the equivalent experimental data (dots). Simulations beyond the range of antibody concentrations used in the experimental data are indicated with dashed lines. Experimental data was not obtained for the combination of tocilizumab and 10H2, but simulations are presented here for comparison. Model output and experimental data are each normalized to the bound BS1 concentration from the initial antibody concentrations where binding reached saturation. The error bars depict the standard error from three experimental replicates; the experimental data was previously published [28]. Simulation results with all obtained parameter sets are included in the *Supplemental Information* [**Figure S7**].

In the single-receptor-positive cell lines (denoted IL-6R^+^ and IL-8R^+^), at low antibody concentrations, there is greater binding of the monospecific antibodies than of BS1 because each monospecific antibody molecule has two binding sites (doubling overall likelihood of binding), plus avidity effects promote increased binding. As the antibody concentration increases within the experimentally-tested range, the monospecific antibody curves appear to saturate at a lower level of total antibody bound than BS1 (consistent with the experimental measurements) because much of the monospecific antibody is bound bivalently, with a single antibody occupying two receptors. In contrast, BS1 can only bind monovalently because its second binding site is for a receptor not expressed on that cell type. Thus, BS1 forms antibody-receptor complexes whereas tocilizumab and 10H2 form receptor-antibody-receptor complexes, resulting in a lower measured signal since binding is quantified by the number of antibodies bound.

At simulated antibody concentrations higher than those experimentally tested [**Figure 3, dashed lines**], we predict further increased binding of the monospecific antibodies; at very high antibody concentrations, the antibody is present in such excess that all of the antibody is bound monovalently, matching the behavior of BS1, i.e., receptor-antibody-receptor complexes are lost in favor of antibody-receptor complexes. As a result, at the highest concentrations, the overall binding of each antibody is predicted to be equivalent; however, at practical experimental concentrations, we can see and explain the higher binding of BS1.

In these single-receptor-positive cell lines, BS1 has a sigmoidal binding curve because it is effectively monovalent; however, in the double-receptor-positive cell line (denoted IL-6R^+^/IL- 8R^+^, which expresses about twice as many IL-8 as IL-6 receptors [**Table 1**] [28]), BS1 is now effectively bivalent and exhibits a binding curve similar to and higher than the individual tocilizumab and 10H2 curves. This shape is due to BS1 bivalently binding both IL-6R and IL-8R simultaneously, and the total BS1 binding at the highest levels [**Figure 3, dashed lines**] is higher than either tocilizumab or 10H2 because it can bind to either IL-6R or IL-8R. The bound receptor concentrations can be further separated by complex type, distinguishing binary antibody-receptor complexes from ternary receptor-antibody-receptor complexes [**Figure S6**]. The separation of complex types confirms that the shape of the binding curve in the double-receptor-positive cell line is due to the formation of ternary complexes through bivalent binding.

We also simulated the exposure of each of the three cell types to a combination of the two monospecific antibodies (tocilizumab + 10H2), and, as expected, in simulations of single-receptor-positive cell lines, the combination behaved similarly to the single monospecific; while in the double-receptor-positive cell lines, the combination behaved similar to the bispecific. This has potentially useful implications for the ability of the bispecific (vs the monospecific combination) to bind to and inhibit receptors on cells expressing these two receptors at different levels.

### Bivalent Antibody Binding over Time

We simulated the formation of the BS1-receptor complexes over time [**Figure 4**], using an equivalent timeline to the experiments: an initial 2-hour binding period, followed by a washout of all free (unbound) BS1 from the system at t = 2 hours to simulate dissociation of the antibody-receptor complexes. The initial antibody concentration was set to 100 nM to ensure the antibody fully saturated the available receptor by 2 hours; similar results are also demonstrated for lower antibody concentrations [**Figure S8A**].

**Figure 4.**
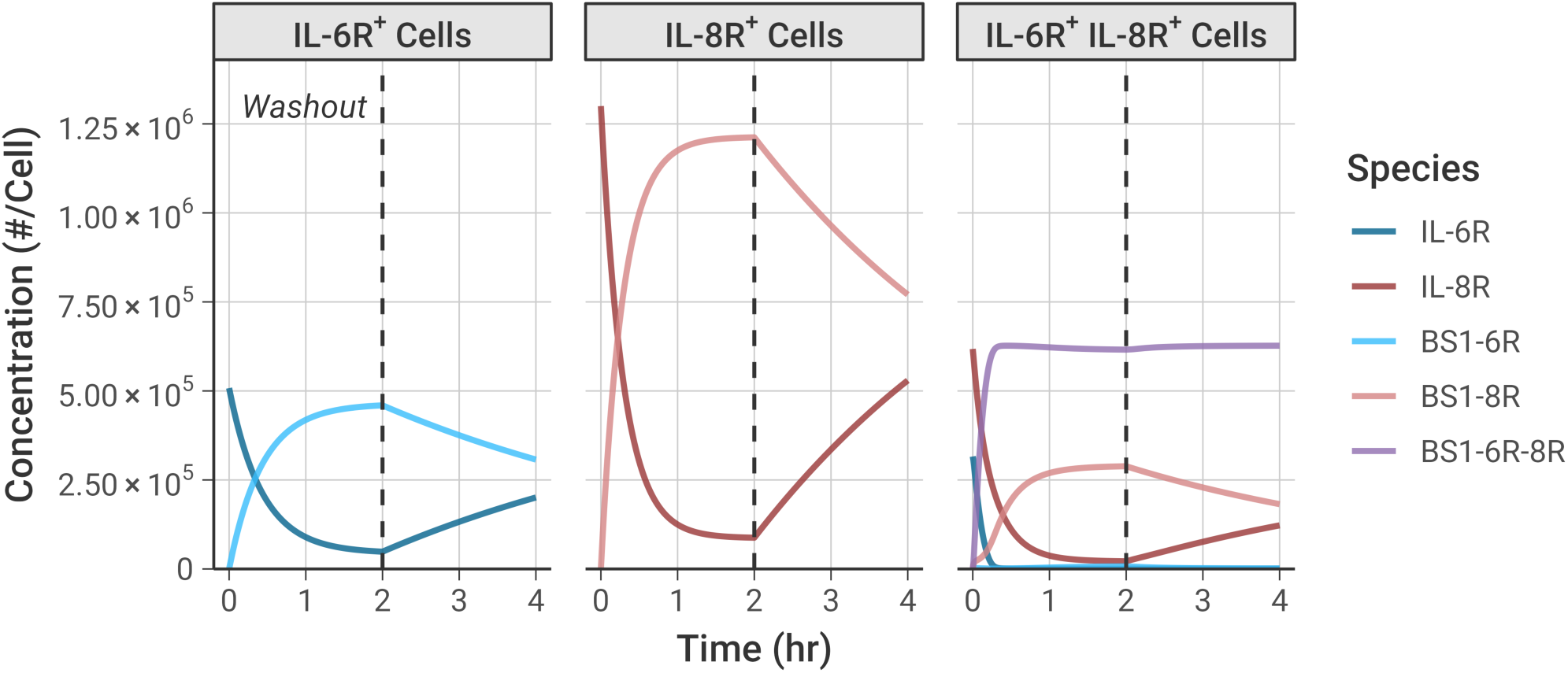
Simulations illustrate the dynamics of BS1 antibody binding to IL-6R and IL-8R over time. Initial BS1 concentration = 100 nM and 10^5^ cells/well for all simulations. Free (unbound) BS1 concentration was set to 0 nM at 2 hours to simulate antibody washout from the system. The expression levels of IL-6R and IL-8R from the transduced experimental cell lines [**Table 1**] were used in the simulations. Simulation results for additional antibodies and antibody concentrations are included in the *Supplemental Information* [**Figure S8**].

Binding of BS1 in the single receptor-positive cell lines, IL-6R^+^ and IL-8R^+^, shows formation of binary antibody-receptor complexes in the association phase, followed by dissociation of those complexes in the washout phase. Binding is greater in the IL-8R^+^ cell line because there is a higher receptor expression in those cells than in the other cell lines [**Table 1**].

In the double-receptor-positive cell line, IL-6R^+^/IL-8R^+^, the association phase shows substantial ternary IL-6R-BS1-IL-8R complex formation [**Figure 4, purple line**], with fewer binary BS1-IL-8R complexes and almost no BS1-IL-6R complexes being formed. Initially, formation of the ternary complexes progresses rapidly, quickly reaching a steady concentration. After the first 20 minutes, as the free IL-6R becomes saturated with antibody, more binary BS1- IL-8R complex formation occurs because there is no free IL-6R remaining to form ternary complexes. A small amount of binary BS1-IL-6R complex forms near the end of the association phase, but there is much greater binary BS1-IL-8R complex formation because IL-8R is in excess of IL-6R in this cell line.

During the dissociation phase in the IL-6R^+^/IL-8R^+^ cells, the concentration of the binary BS1-IL-6R and BS1-IL-8R complexes decreases as antibody unbinds due to mass action following removal of the excess free (unbound) antibody. Perhaps counterintuitively, the concentration of the ternary IL-6R-BS1-IL-8R complexes actually slightly increases in this phase, as more receptors are freed and become available to bind to existing binary complexes, and the ternary complex concentration achieves a steady value and does not decrease during the simulation time period. This same behavior is also observed in binding of the two monospecific antibodies, tocilizumab and 10H2, to their target receptors [**Figures S8B, S8C**]. This illustrates the importance of avidity in bivalent antibody binding; each of the antibodies is able to form ternary complexes as more receptor becomes available for binding, and these complexes persist at a consistent concentration for a significant period of time after antibody removal.

### Effect of Varying Antibody Concentration and Receptor Expression

To better understand the impact of the antibody concentration on bispecific antibody-receptor complex formation, and in particular on the relative formation of binary (BS1-IL-6R or BS1-IL- 8R) versus ternary (IL-6R-BS1-IL-8R) complexes, we performed simulations of BS1 binding over a range of initial antibody concentrations and for cells with differing levels of combined IL-6R and IL-8R expression [**Figure 5**]. Because BS1 requires both IL-6R and IL-8R available to form ternary complexes, BS1-receptor complex formation is very sensitive to the ratio of IL-6R to IL- 8R expression in the system. Thus, to specifically isolate the impact of overall receptor expression on BS1 binding, IL-6R and IL-8R were kept in a 1:1 ratio for these simulations. In these results and the results that follow, the bound receptor is reported as the receptor fractional occupancy, which is calculated as the fraction of the total receptor concentration (IL-6R + IL-8R) that is bound in a particular complex type (i.e., binary or ternary) or that is bound overall (i.e., total bound).

**Figure 5.**
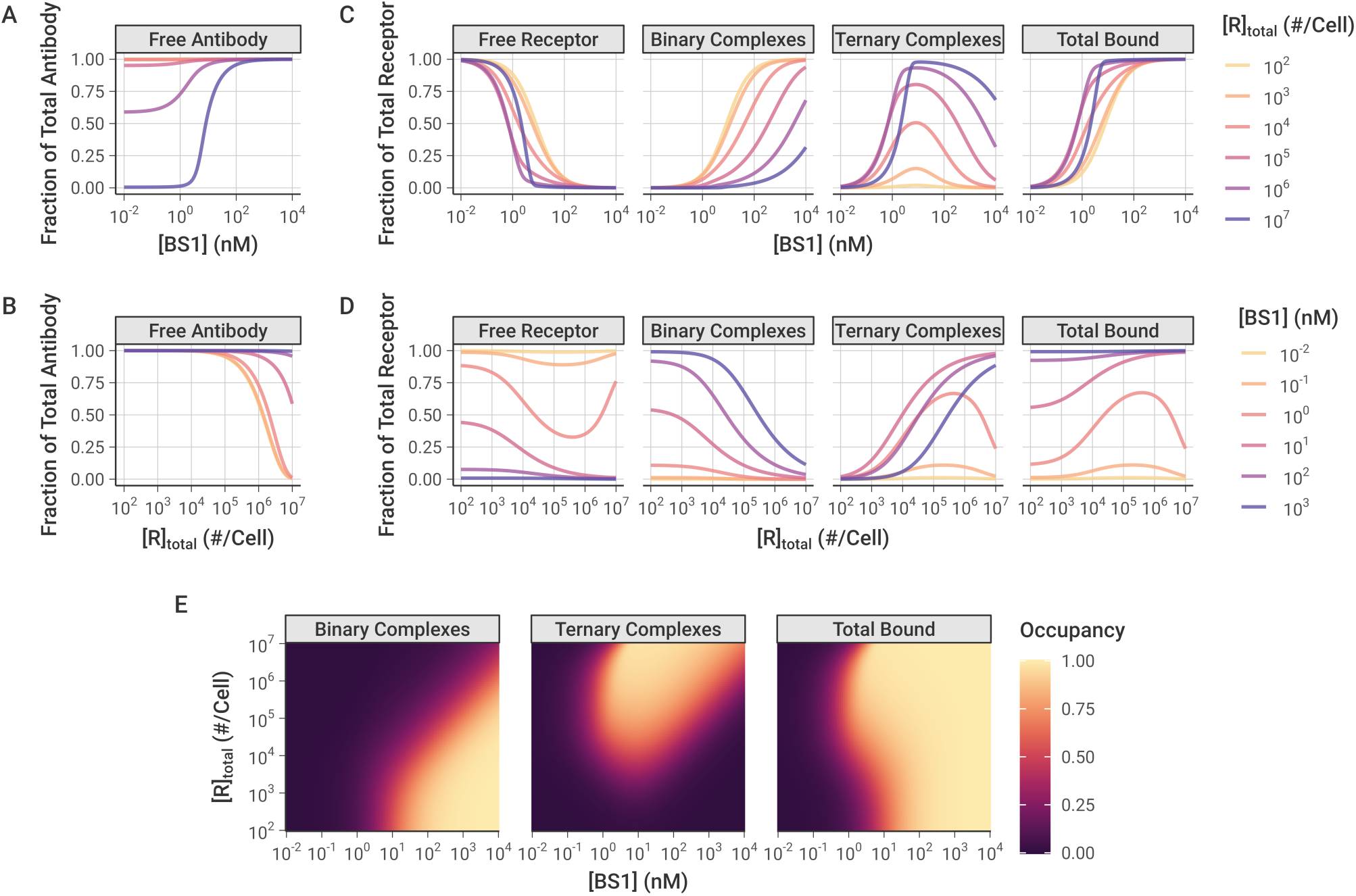
Simulated Binary (Ab-R), Ternary (R-Ab-R), and Total Bound (Binary + Ternary) levels of bispecific BS1-Receptor complexes, over a range of antibody doses and receptor expression levels. In these simulations, IL-6R and IL-8R are present in a 1:1 ratio, and simulations were performed for 24 hours after antibody dosing. **A-B,** Fraction of total BS1 concentration in free (unbound) state for different levels of initial BS1 (**A**) and receptor (**B**). **C-D,** Fraction of total receptor (IL-6R + IL-8R) in different forms/complexes for different levels of initial BS1 (**C**) and receptor (**D**). **E,** Bound receptor fraction across different levels of initial BS1 and receptors. The color indicates the fraction of the total receptor (IL-6R + IL-8R) that is bound to BS1 in each antibody-receptor complex type. A similar heat map for different tocilizumab and 10H2 concentrations is included in the *Supplemental Information* [**Figure S9**].

At higher antibody concentrations and lower receptor expression levels, the antibody levels are saturating and very little of the antibody is consumed in the bound complexes [**Figures 5A, 5B**]. At lower antibody concentrations and higher receptor expression levels, however, the free receptor is in excess of the free antibody and a substantial fraction of the antibody is bound to receptor (up to 100 percent at the highest simulated receptor expression level).

The relative excess of antibody or receptor is important for the proportion of binary antibody-receptor and ternary receptor-antibody-receptor complexes that are formed. Ternary complexes require two dissociation reactions to fully separate, and binary complexes tether the antibodies in close proximity to free receptor, promoting recreation of ternary complexes when they do dissociate; thus, ternary complexes represent a more stable antibody binding format relative to binary complexes. When the antibody is present in excess of the receptor (i.e., at high antibody concentrations and low receptor expression), the majority of complexes that form are the less-stable binary complexes. Additionally, at any given receptor density, as the concentration of antibody is increased, more binary complexes are created [**Figures 5C, 5D**].

Ternary receptor-antibody-receptor complexes, in contrast, are favored at intermediate antibody concentrations, around 10^0^ to 10^2^ nM (recall **Figure 3**). Initially, increasing the antibody concentration causes more ternary complexes to form, but, past a certain threshold, the free antibody overwhelms the number of available receptors. At these higher antibody concentrations, there are few remaining receptors available for the second binding reaction to convert binary complexes into ternary complexes [**Figures 5C, 5D**]. This bell-shaped relationship between ternary systems and bivalent molecule concentration has been described previously [53]; the decreased ternary complex formation at high concentrations is termed “autoinhibition.”

Interestingly, model simulations demonstrate that, at a constant antibody concentration, as the receptor expression is increased, a greater fraction of the receptor is bound to antibody [**Figure 5D**]. This initially appears counterintuitive because increasing receptor expression means the system contains more binding sites for the same amount of antibody. However, this result can be rationalized by the fact that the increase in receptor expression causes a greater proportion of the bound complexes to be of the higher-stability ternary format, which benefit from avidity effects and therefore readily rebind if one arm dissociates, making them less likely to fully dissociate.

Increased proportion of bound receptor with increasing receptor concentration is only observed when the antibody amount is still in excess of the receptor. At higher receptor expression levels with lower antibody concentrations, the receptor becomes the excess molecule, resulting in a greater proportion of the receptor remaining unbound [**Figure 5D**].

The combined impact of antibody concentration and receptor expression creates “zones”, wherein different antibody-receptor complex types are favored [**Figure 5E**]. Overall, increasing the antibody concentration causes more of the receptor to be bound in total. Binary antibody-receptor complexes are favored at higher antibody concentrations and lower receptor expression levels; whereas, ternary receptor-antibody-receptor complexes are the predominant type at high receptor expression levels with intermediate to high antibody concentrations. This same pattern is also observed in simulations of combination treatment with the two monospecific antibodies, tocilizumab and 10H2, modeled together in a 1:1 concentration ratio [**Figure S9**].

### Monovalent and Bivalent Binding

To further explore how ternary receptor-antibody-receptor complex formation leads to greater fractional receptor binding as receptor expression increases, we compared the bivalent antibody binding behavior to simulations of BS1 that we artificially restricted to monovalent binding only [**Figure 6**]. In these simulations, the association rate constants for the second binding step (kon,6R* and kon,8R*) were fixed at 0 to prevent ternary complex formation and restrict BS1 to monovalent binding only. The rate constants for the initial association into binary complexes (kon,6R and kon,8R) and for the dissociation of antibody-receptor complexes (koff,6R and koff,8R) were kept at their previous values [**Table 2**].

**Figure 6.**
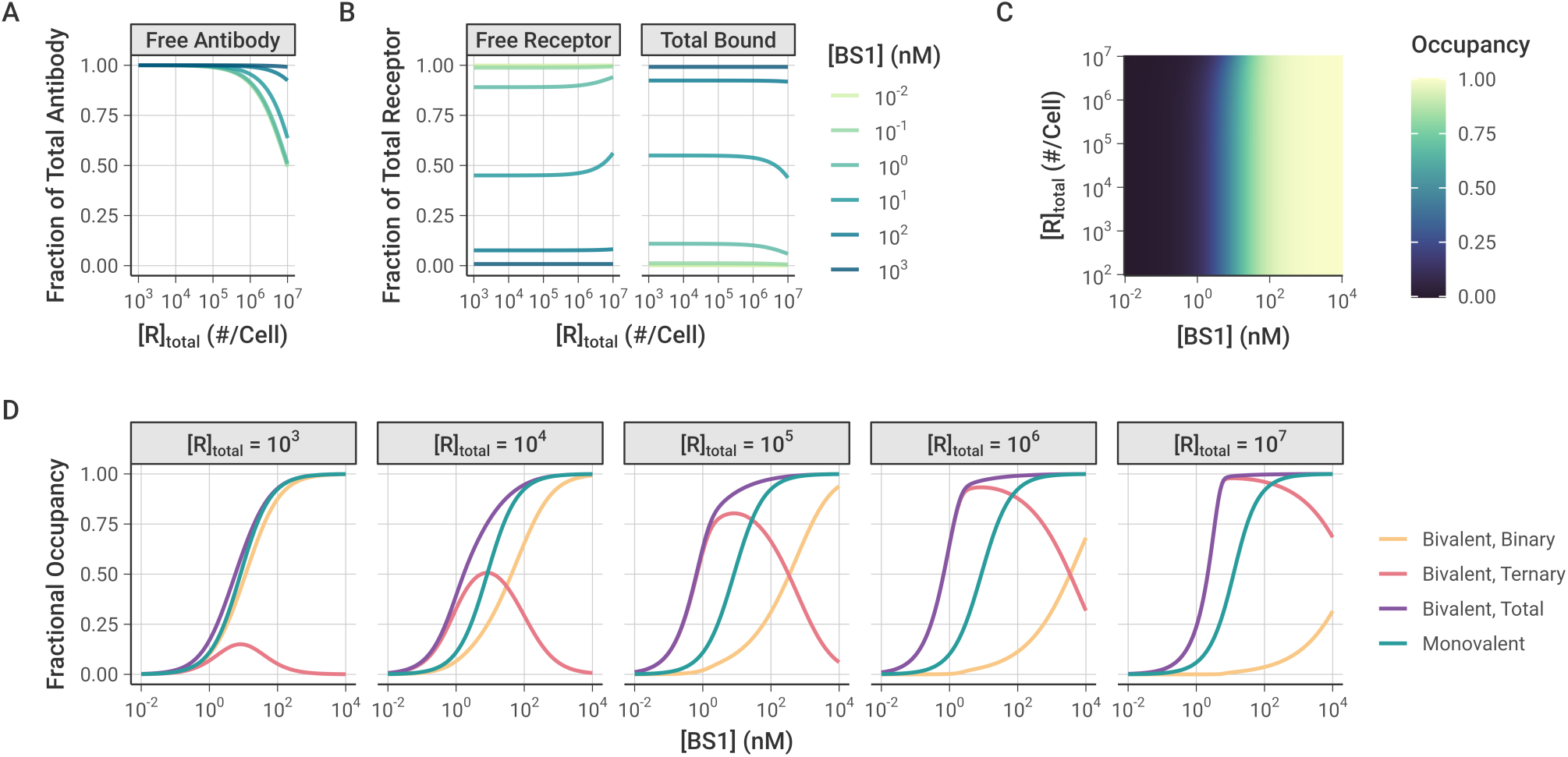
Simulations of monovalent BS1 binding over varying initial antibody and receptor concentrations. In these simulations, although the cells express both receptors, the formation of ternary complexes was suppressed by setting kon,6R* and kon,8R*to 0. IL-6R and IL-8R are present in a 1:1 ratio, and simulations were performed for 24 hours after antibody dosing. Similar results for the combination of the monospecific antibodies tocilizumab and 10H2 are included in the *Supplemental Information* [**Figure S11**]. **A,** Fraction of total BS1 that is free (unbound) for different levels of receptor expression and initial BS1 concentration. **B,** Fraction of total receptor concentration (IL-6R + IL-8R) that is unbound (free) or bound (in binary antibody-receptor complexes) for different levels of receptor expression and initial BS1 concentration. The same results, but with antibody and receptor visualization reversed, are included in the *Supplemental Information* [**Figure S10**]. **C,** Bound receptor fraction across different initial BS1 and receptor levels. The color indicates the fraction of the total receptor (IL-6R + IL-8R) that is bound to antibody. **D,** Comparison of monovalent and bivalent binding. The lines indicate the fraction of total receptor (IL-6R + IL-8R) that is bound in different complex types in the original simulations and the simulations restricted to monovalent binding only. Each panel represents a different receptor level (in # receptors/cell).

Similar to the simulations of bivalent BS1 binding, at lower receptor expression levels, the antibody is present in excess of the receptor and fully saturates the receptor [**Figures 6A, S10**]. At the highest receptor expression levels, more of the antibody is consumed in the binding, but there is still a substantial fraction of free antibody available. Unlike the previous bivalent binding simulations, however, the simulations of monovalent binding show that the fraction of receptor that is bound to antibody is nearly entirely independent of the receptor expression. For most of the receptor levels, the bound receptor fraction varies only with the initial antibody concentration and remains constant as receptor expression is increased [**Figures 6B, 6C**]. The only deviation from this pattern is at the highest receptor expression levels for intermediate antibody concentrations, wherein the antibody is no longer saturating the receptor, leading to a greater proportion of free receptor remaining.

When the antibody is restricted to only binding monovalently, as long as the antibody is present in excess of the receptor, varying the receptor expression level does not change the proportion of the receptor that is bound. In contrast, when the antibody binds bivalently, the receptor expression level in the system determines the proportion of binary and ternary complexes that form [**Figure 6D**]. At the lower receptor expression levels, the antibody fully saturates the receptor and the receptor is primarily bound in binary antibody-receptor complexes. As the receptor expression increases, the proportion of receptor in binary complexes decreases and ternary complexes begin to dominate. Because ternary complexes are the more stable form, the proportion of bound receptor overall also increases, leading to the previously illustrated pattern of greater fractional receptor binding with increasing receptor expression for a given antibody concentration. While these results focus on the bispecific antibody BS1, this is a general pattern of bivalent antibody binding and is seen for the monospecific antibodies tocilizumab and 10H2 as well [**Figure S11**].

### Comparison between Monospecific and Bispecific Antibodies

Tocilizumab, 10H2, and BS1 are all bivalent antibodies that can form both binary antibody-receptor and ternary receptor-antibody-receptor complexes when exposed to their respective target receptors. Thus, they all share the previously discussed binding behaviors across varying total antibody and receptor concentrations. However, BS1 differs from tocilizumab and 10H2 in that it is bispecific and simultaneously binds to both IL-6R and IL-8R. To further examine how antibody-receptor complex formation compares between monospecific and bispecific antibodies, we simulated BS1 and the combination of tocilizumab and 10H2 over a range of different IL-6R and IL-8R expression levels [**Figure 7A**]. In these simulations, the total initial antibody concentration was held constant at 10 nM, and tocilizumab and 10H2 were combined in a 1:1 ratio. The combination of tocilizumab and 10H2 targets both IL-6R and IL-8R but differs from BS1 in that each antibody can only bind one type of receptor. Although earlier results were presented for an initial antibody concentration of 100 nM [**Figure 4**], the overall binding is strong at 100 nM and the differences between the antibody types are less apparent. Results for additional antibody concentrations are included in the *Supplemental Information* [**Figure S12**].

**Figure 7.**
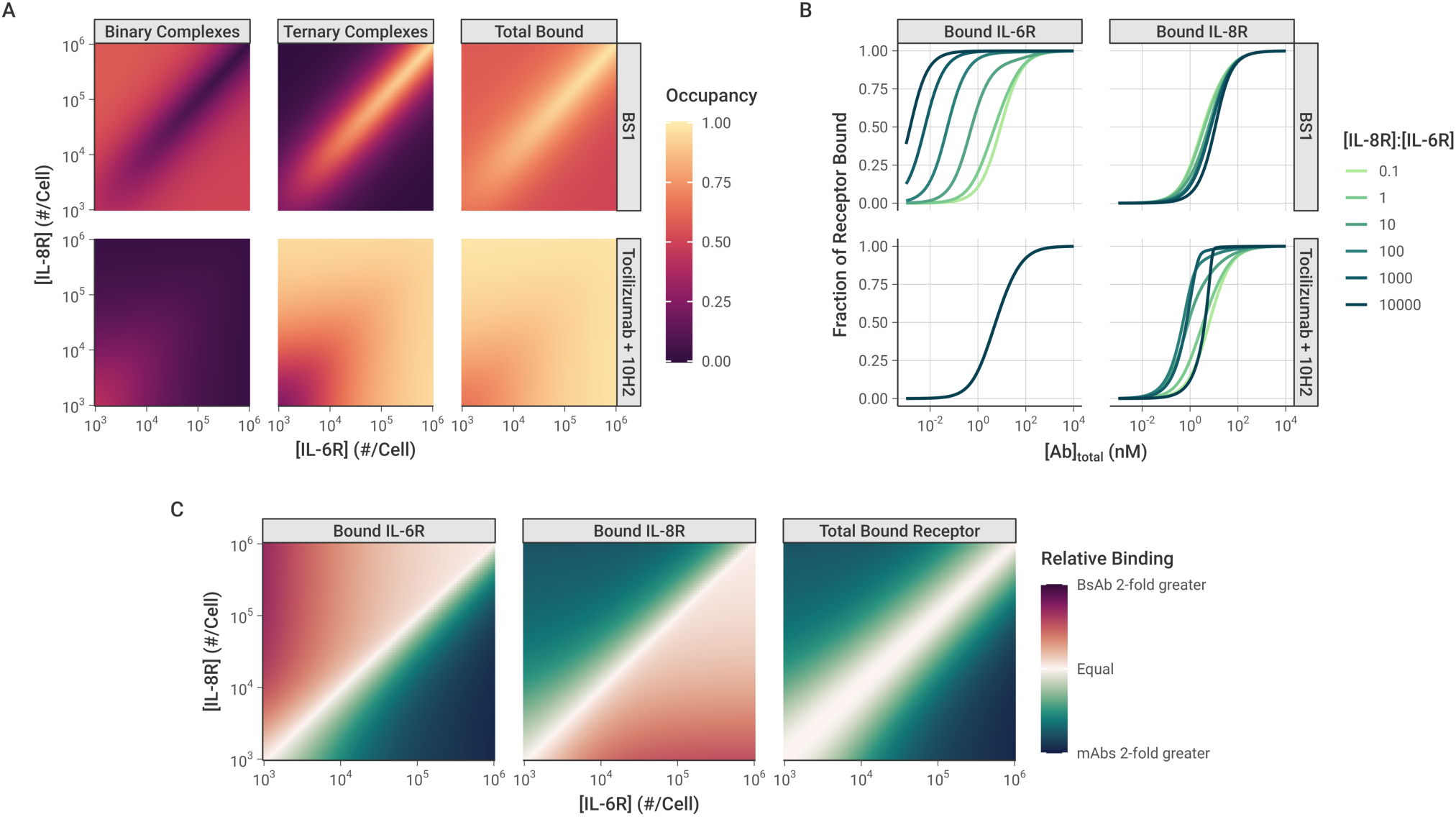
Comparison of antibody-receptor complex formation: BS1 vs. combination of tocilizumab and 10H2. All simulations were performed for 24 hours after antibody dosing. **A,** Fraction of all receptors (IL- 6R + IL-8R) that are bound in Binary and Ternary complexes, and Total Bound receptor (Binary + Ternary) across different IL-6R and IL-8R concentrations. The color indicates the fraction of all receptors (IL-6R + IL- 8R) that are bound in each antibody-receptor complex type. Initial BS1 concentration = 10 nM; initial tocilizumab concentration = 5 nM and initial 10H2 concentration = 5 nM. Heat maps of additional total antibody concentrations are available in the *Supplemental Information* [**Figure S12**]. **B,** The fractional occupancy of each receptor individually when one receptor (IL-8R) is in excess. IL-6R was fixed at 10^3^ receptors/cell for these simulations, while IL-8R ranged from 10^2^ to 10^7^ receptors/cell. The fractional occupancy indicates the fraction of the specific receptor concentration (either IL-6R or IL-8R) that is bound to antibody (either BS1 or the combination of tocilizumab and 10H2). Results with IL-8R as the fixed receptor are included in the *Supplemental Information* [**Figure S13**]. **C,** Comparison of receptor bound by BS1 (BsAb) or the combination of tocilizumab and 10H2 (mAbs) across different receptor concentrations. Relative binding is the ratio of fractional bound receptor (the fraction of total IL-6R + IL-8R bound to antibody) when BS1 is used compared to when the combination of mAbs is used. Similar heat maps for different total antibody concentrations are included in the *Supplemental Information* [**Figure S14**].

As was demonstrated previously, when the receptors are present in a 1:1 ratio, the complex formation is identical between BS1 and the combination of monospecific antibodies [**Figures 5E, S9**], and this behavior holds across antibody concentrations [**Figure S12**]. Outside of a 1:1 IL-6R:IL-8R ratio, however, the antibodies demonstrate very different binding behaviors. BS1 requires both IL-6R and IL-8R to be available to form ternary complexes, so BS1 ternary complex formation is favored when the receptors are present in a 1:1 ratio [**Figure 7A**]. When either receptor is present in excess of the other, ternary complex formation is limited by the lower-expressed receptor. In this case, the excess receptor will only be able to bind to BS1 in a less-stable binary complex, leading to lower binding overall.

In contrast, because tocilizumab and 10H2 each bind to one type of receptor, the ratio of IL-6R to IL-8R expression does not impact their binding; only the total receptor expression has an effect on antibody-receptor complex formation for the monospecific antibodies [**Figure 7A**]. At lower receptor levels where the antibodies are present in substantial excess over the receptors, binary complexes are favored. As the total receptor expression increases, more receptors are available to form ternary complexes, and complex binding shifts to favor the ternary form.

Similar trends are observed at different total antibody concentrations as well [**Figure S12**]. At lower antibody concentrations, there is less complex formation overall for all antibodies. Notably, at the highest receptor levels, the receptor is present in excess of the antibody, leading to a low fractional occupancy of the receptor. In comparison, higher antibody concentrations lead to greater antibody-receptor complex formation across all receptor expression levels, but otherwise show the same pattern of binding behavior.

When the complex formation is examined for each receptor separately, however, a distinct pattern emerges [**Figure 7B**]. In these simulations, the concentration of IL-6R was held constant at 10^3^ receptors per cell, while the concentration of IL-8R ranged from 10^2^ to 10^7^ receptors per cell. The proportion of each individual receptor that is bound is reported. IL-8R was simulated as the excess receptor because it has been shown to be up-regulated relative to IL-6R in breast cancer [28], and similar results are shown when for simulations with IL-6R in excess [**Figure S13**]. The monospecific antibodies (tocilizumab and 10H2) each bind to a single receptor type, so the binding of each receptor is independent. Thus, for the combination treatment, varying IL-8R concentration has no effect on the amount of bound IL-6R, and the bound IL-8R concentration shows identical behavior to the simulations where the total receptor concentration was varied [**Figures 7B, S9**].

The binding of BS1, however, is highly dependent on the ratio of IL-6R to IL-8R expression. As the concentration of IL-8R (the excess receptor in this case) increases, BS1 shows increasing occupancy of IL-6R, the limited receptor [**Figure 7B**]. In these simulations, the concentration of IL-6R was not varied, so the change in IL-6R binding is driven entirely by the increased IL-8R concentration. As greater IL-8R is present in the system, BS1 forms more binary BS1-IL-8R complexes, tethering it to the cell surface and bringing it within close proximity of the free IL-6R. This increases BS1 binding to IL-6R, even when it is present at substantially lower concentrations than the other receptor. This behavior is observed only for the bispecific antibody and is driven by the initial interaction with the excess receptor. The tradeoff, however, is that BS1 shows lower occupancy of the excess receptor, IL-8R, at intermediate antibody concentrations, compared to the combination of monospecific antibodies, and the occupancy declines as the IL-8R concentration increases and the ratio of IL-6R to IL-8R moves further away from 1:1.

The impact of these differing binding patterns is apparent when the monospecific and bispecific antibodies are compared directly [**Figure 7C**]. The relative binding output from these simulations quantifies the fold-change in fractional occupancy of each receptor when BS1 binds compared to the binding of the combination of tocilizumab and 10H2. Similar to the previous results, for the overall bound receptor, the monospecific and bispecific antibodies show the same results when the receptors are present at a 1:1 ratio, and the monospecific antibodies show greater binding outside of this ratio. However, the occupancy of the individual receptors reveals that, when one receptor is present in excess of the other, the monospecific antibodies show greater binding to the excess receptor, while BS1 binds more of the limited receptor. The fold-change in binding between the antibody types increases as the ratio moves further away from equal receptor expression [**Figure 7C**]. This pattern is observed for additional antibody concentrations as well, but the differences between antibody types diminish as the concentration increases because the overall binding is high [**Figure S14**]. Overall, these results suggest the binding of the bispecific antibody (but not monospecific antibodies) to a low concentration receptor is enriched by the presence of a high concentration of the other target molecule.

### Univariate Sensitivity

To analyze the impact of individual model parameters on the bispecific antibody binding, we performed local and global univariate sensitivity analyses of the model output [**Figure 8**]. First, we examined the local sensitivity of ternary IL-6R-BS1-IL-8R complex formation and total receptor binding to changes in the values of the association and dissociation rate constants along with initial antibody and receptor concentrations [**Figure 8A**].

**Figure 8.**
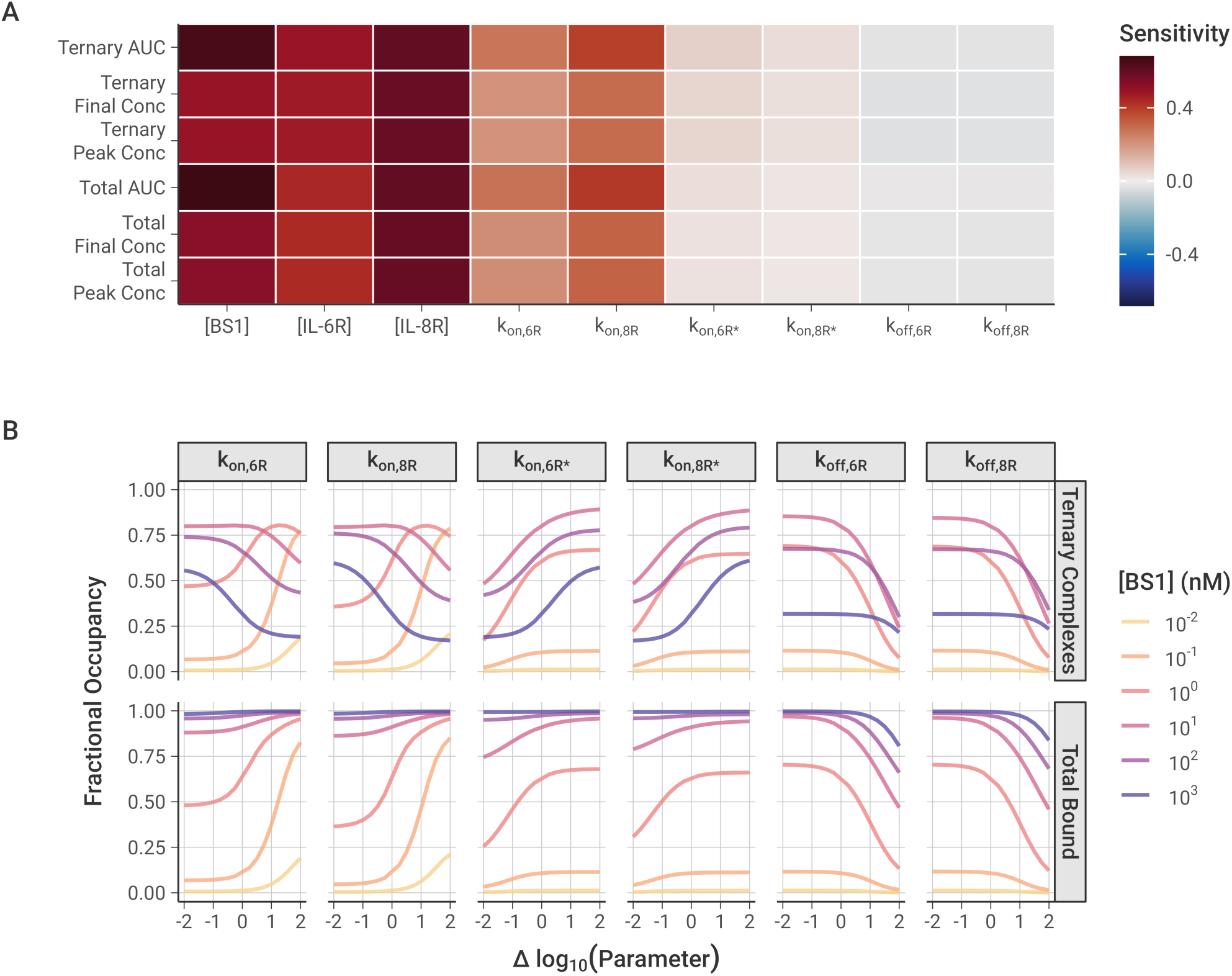
Local and global sensitivity of model output to association and dissociation rate constants and the initial antibody and receptor concentrations. **A,** Local sensitivity analysis of model output with varying rate constants and initial concentrations. 10 nM baseline BS1 concentration, [IL-6R] = [IL-8R] = 5 × 10^4^ receptors/cell, and output at t = 2 hours for all simulations. Area Under the Curve (AUC) is calculated as the integration of the BS1-receptor complex concentration over time, determined for the ternary complexes and for the total bound receptor (IL-6R + IL-8R). Sensitivity is calculated as the percentage change in the output divided by the percentage change in the parameter (10% for these simulations). **B,** Global sensitivity of fractional bound receptor concentration over varying rate constant value. Each parameter was varied over two orders of magnitude below and above its optimized value. Fractional occupancy is determined as the fraction of total receptor (IL-6R + IL-8R) that is bound to BS1 in a particular complex type, separated for ternary complexes and total bound in binary or ternary complexes. Simulations were performed for 24 hours after antibody dosing, and [IL-6R] = [IL-8R] = 5 × 10^4^ receptors/cell for all simulations.

Overall, the antibody-receptor complex formation is more sensitive to the initial antibody and receptor concentrations than it is to the binding rates, with the BS1 and IL-8R concentrations having the greatest impact on binding. At two hours after antibody dosing, the amount of bound complex is still increasing [**Figures 4, S8**], so increasing the concentration any of the molecules in the system will lead to greater binding. The concentration of IL-8R is slightly more impactful than that of IL-6R because the binding to IL-8R is estimated here to be slightly faster than the binding to IL-6R [**Table 2**], leading to more binary complexes and then more ternary complexes being formed.

Of the association and dissociation rate constants, the model output is most sensitive to the initial binding step to form binary complexes, which is the rate-limiting step. The rate of the second receptor binding leading to ternary complexes is so fast that the binding is able to progress immediately after binary complexes are formed, and it does not have a significant impact on the total amount of bound receptor. Increasing the dissociation rates has a very slight negative effect on the receptor binding, but the rates are so slow that little dissociation occurs during the two-hour simulation period and the impact of varying the rate constants is minimal.

With global sensitivity analysis of the model parameters, we examined the impact of varying the rate constant values over a wider range [**Figure 8B**]. The results similarly demonstrate that the total bound receptor is most sensitive to increasing the rate of the first binding step. This is especially true for the intermediate antibody concentrations where neither the antibody nor the receptor are significantly in excess. Decreasing either of the first binding step rate constants from their baseline (i.e., moving left of zero on the x-axis for kon,6R or kon,8R) individually does not have a large impact on the total receptor binding because the antibody can still bind to the opposite receptor to form binary complexes.

Some of those binary complexes may progress forward to forming ternary complexes [**Figure 1C**], but the ternary complex concentration shows very distinct behaviors depending on the antibody concentration. For lower antibody concentrations, increasing the rate of the first binding step leads to more ternary binding, as the limiting factor is the binary complex concentration. For higher antibody concentrations, however, the opposite effect is observed: increasing the rate of binary binding leads to fewer ternary complexes, due to auto-inhibition with the binary complexes consuming all of the available receptor. This is consistent with “zones” of different dominant complex types when the concentrations are varied [**Figure 5E**] and suggests bispecific antibody binding to form ternary complexes is dependent on the balance of binding site affinity and species concentrations.

For the second binding step leading to the ternary complex, the concentrations of all complexes are sensitive to decreasing the rate constant from the baseline but less so to increasing it [**Figure 8B**]. The formation of ternary complexes progresses incredibly quickly relative to the binary complex binding, so increasing their binding rate has little effect. However, slowing the rate of ternary complex binding causes more receptor to be consumed by less-stable binary form, leading to less receptor being available for ternary complex binding and less bound receptor overall. Finally, the amount of bound receptor is only sensitive to increasing the dissociation rate at high magnitudes [**Figure 8B**]. Generally, the dissociation is so slow relative to the association that the specific value does not have a significant impact on the receptor binding, but, at very high rates, enough dissociation occurs that it decreases the number of bound complexes that are present. These results agree with the results from the local sensitivity analysis, collectively demonstrating that the first binding step is the rate-limiting step and that only extreme values of the second binding step and the dissociation rates have a substantial impact on receptor binding.

## DISCUSSION

In this study, we have developed a model of a bispecific antibody (BS1) targeting two key cell surface receptors, IL-6Rα and IL-8RB, which were recently implicated in a synergistic pathway that drives tumor metastasis [12,28]. Our model is comprised of a series of ODEs for each of the receptors, antibodies, and antibody-receptor complexes studied. Each association and dissociation process in the system is represented as a set of terms in the ODEs, with separate terms for the formation of binary antibody-receptor and ternary receptor-antibody-receptor complexes. Related approaches have been used in other mechanistic models of bivalent antibody binding [24,38,39], and below we describe the differences between those works and this. We used *in vitro* experimental data to estimate values for the binding parameters of the model; we observed that the simulations match the experimental results and the data constrains the parameters well. Deploying the model to simulate antibody binding to cells that express one or other or both of the target receptors, and comparing to simulations of combinations of monospecific antibodies, we gleaned insights into the mechanistic differences between these potential treatments.

Once one arm of the antibody binds a cell-surface receptor, the second receptor must be within reach to allow for a second binding event. The distance between antigen-binding domains of IgG antibodies generally ranges from 6 to 12 nm [54–56], but the arms are joined by a highly flexible hinge region that can allow the arms to reach up to 17 to 18 nm when fully extended [57]. Assuming uniform receptor distribution and a surface area of 1000 μm^2^, there would be an average distance between receptors of around 50 nm for 10^5^ receptors/cell or 500 nm for 10^3^ receptors/cell. However, bispecific antibodies have been demonstrated to simultaneously engage two receptors at these receptor densities [33,58]. It has been hypothesized that this is due to fast diffusion of receptors in the cell membrane [40,59] and non-uniform receptor distribution, with receptors being co-localized in lipid rafts and other membrane structures, increasing local density [60,61].

Binding of one arm of the antibody increases the local concentration of the antibody at the surface, leading to a significantly stronger apparent affinity for second arm binding (compared to first arm binding) to form the ternary complex [30,50]. This may be partly reduced by a loss in rotational flexibility and by steric hindrance from other antibody-receptor complexes in the vicinity [37,62], but, on balance, we expect the second binding event to be effectively stronger than the first.

Previous mechanistic bivalent binding models have incorporated the effects of binding avidity in various ways. The models presented by van Steeg et al. [24] and Rhoden et al. [39] use the same association rates for both binary and ternary complex formation, but they instead use the effective local antigen concentration within reach of the bound antibody in the rate equation for the second binding event. Flexibility limitations and steric constraints are not explicitly included in these models. Vauquelin and Charlton [37] likewise scaled the ternary complex formation by the effective local concentration, but they also incorporate a “penalty factor” to the second association constant to account for the limited rotational freedom of the bound molecule. Finally, Harms et al. [38] incorporated both the heightened effective concentration and the restricted flexibility for bivalent binding into a single parameter they termed as the “cross-arm binding efficiency” (χ). They defined χ as the ratio of the kon for ternary complex formation to the kon for the initial binary complex binding (represented in our model as kon,R* and kon,R, respectively) and hypothesized that χ was an epitope-dependent property of a bivalent antibody, independent of the antibody’s monovalent affinity for its target [51].

Since we had an experimental data set for our system, we did not need to make assumptions about the relative strength of first versus second binding, and instead estimated independent values for these steps using empirical data. The ratio between these first and second binding rate constants is the χ parameter defined by Harms et al. [38], and the value we obtained (1.6 × 10^4^) was consistent with the range of values for χ they reported (10^2^ to 10^5^). Larger values indicate stronger cross-arm binding, consistent with our results demonstrating that BS1 binds bivalently to IL-6R and IL-8R with high avidity [**Figures 3, 5A, 5C**].

For our model, several key assumptions made it possible to reduce the number of unknown parameters. First, because BS1 was constructed using the binding domains from tocilizumab and 10H2, we assumed that the antibodies share rate constants for the first binding step to the same receptor. Second, since all three antibodies are bivalent and IgG-based, we assumed the geometries were similar enough that they share rate constants for the second binding step to the same receptor (but note that the first and second step rate constants are different). Third, we assumed that the dissociation rate constants were shared for unbinding events from a specific receptor. The rate constant values are further constrained by detailed balance (see *Methods*). These assumptions are supported by the strong fit of the model to experimental data [**Figures 3, S7**], and the well-constrained parameter sets [**Figures 2, S4**]. If the assumptions were relaxed, the values of the parameters would be less constrained and thus more uncertain.

Simulations of the formation of antibody-receptor complexes over time revealed the binding dynamics in the system [**Figures 4, S8**]. Initially, binary antibody-receptor complex formation progresses quickly, followed by conversion of binary complexes into ternary receptor-antibody-receptor complexes shortly thereafter. When one receptor is in excess of the other, as IL-8R was in our IL-6R^+^/IL-8R^+^ experimental cell line, we found that binary complexes with the excess receptor will continue to accumulate after the ternary complex concentration reaches steady state due to the consumption of the limited receptor. Perhaps surprisingly, after the free antibody concentration is washed out of the system, the ternary complex concentration continues to increase, as binary complexes dissociate and free receptor becomes available for binding again. Increased ternary complex formation as the free antibody concentration declines demonstrates the impact of avidity in bivalent antibody binding – in the single-receptor-positive cell lines where only one domain is capable of binding, there is no increase in bound receptor after the washout phase [**Figure 4**]. This “washout” simulates the effects of decreasing antibody concentration, as might be seen physiologically in the window between therapeutic doses as antibody is cleared, internalized, and degraded. When ternary complexes first dissociate, the antibody remains tethered to the cell surface through its remaining receptor bond, promoting rebinding of the antibody and increasing antibody residence time [31,37].

We also demonstrated that the combined effects of antibody concentration and receptor expression level determine the relative balance of binary and ternary complex formation [**Figure 5**], creating different “zones” wherein different complex types dominate. As ternary complexes are thought to be the key pharmacologically relevant species [21,26,32], elucidating the mechanisms underlying complex type distribution is important for successful bispecific targeting. Our simulations show that when antibody (either monospecific or bispecific) is present in excess of the receptor concentration, the less-stable binary complex form is favored; whereas the ternary complex form is dominant in the window of intermediate antibody concentrations and higher receptor expression levels [**Figure 5E**]. This result suggests an optimal therapeutic window for the bispecific antibody therapeutic where maximal binding can be achieved.

Intriguingly, our results also revealed that as surface receptor expression levels increased for a given antibody concentration (monospecific or bispecific), the fractional occupancy of the receptor also increased [**Figures 5B, 5D, S9**]. At first, this behavior seems counterintuitive, since the system is gaining more binding sites for the same number of antibody molecules. To clarify why this pattern appears, we simulated BS1 binding compared to the binding of a theoretical “monovalent” antibody that was restricted to forming only binary complexes [**Figure 6**]. Over the majority of the receptor concentration range tested, the monovalent-restricted antibody binding is independent of the varying receptor level [**Figure 6B**], indicating that it is specifically the formation of ternary complexes that drives the increased receptor occupancy at higher receptor concentrations [**Figure 6D**].

Sensitivity analysis of model binding parameters showed that maximal formation of ternary complexes depends on the balance of antibody concentration and the rates of initial association to form the binary antibody-receptor complexes [**Figure 8B**]. Ternary complex formation by lower affinity antibodies is concentration-limited, with increasing antibody concentration leading to more ternary complex binding. Higher affinity antibodies, however, show “auto-inhibition”, where increasing the amount of antibody leads to the receptor getting overwhelmed by binary complexes, with no free binding sites remaining available for the second association. Ternary binding by these higher affinity antibodies benefits more than ternary binding by lower affinity antibodies from an increased rate of conversion of binary complexes to ternary complexes, which defines the “cross-arm binding efficiency”.

Overall, our results suggest that the binding of bivalent (both monospecific and bispecific) antibodies depends on the interplay of the antibody’s inherent affinity for the target and its cross-linking efficiency, along with both antibody concentration and receptor expression level. There appears to be a “Goldilocks” effect, wherein binding is maximized when affinity, cross-linking, and concentration are all balanced, with none too high or too low. This “affinity and avidity window” has been hypothesized to drive the *in vivo* selectivity of antibodies for target tissues [23,33,63]. Antibodies with intermediate affinity rely on bivalent interactions for stable binding, as opposed to high affinity antibodies that show a greater predilection for monovalent interactions. The requirement for intermediate-affinity antibodies to utilize bivalent interactions may drive selective binding to tissues with greater expression of the target antigen, leading to fewer off-target toxic side effects [63,64]. This is particularly important in cancer, as many of the treatment targets are molecules that are up-regulated in cancerous cells but are still expressed on healthy tissues. These results also suggest a benefit for developing “affinity-modulated” bispecific antibodies, with lower inherent affinities, in order to maximize treatment selectivity. Computational bivalent binding models based on the framework presented here can be applied to specific therapeutic targets to predict optimal affinities (including relative affinities between different binding arms) and cross-linking capacities for maximal binding.

While the total level of receptor expression is a key determinant of bivalent antibody binding, the relative amount of the two different target receptors is also critical for bispecific antibodies. Our simulations revealed that when IL-6R and IL-8R are present in a 1:1 ratio, bispecific antibody binding is nearly identical to the binding of the combination of anti-IL-6R and anti-IL-8R monospecific antibodies [**Figure 7A**]. However, when one receptor is present in excess of the other on the same cell, BS1 shows significantly different behavior at sub-saturating concentrations. Increasing the concentration of the receptor in excess leads to BS1 forming more binary complexes with that receptor. This in turn tethers BS1 to the cell surface, allowing it to rapidly cross-link with the limiting receptor to form the stable ternary complex. The bispecific antibody demonstrates a heightened apparent affinity for the less expressed receptor that increases with the excess receptor concentration [**Figure 7B**]. The monospecific antibodies do not have this advantage; each monospecific antibody has an independent target, and varying expression of one target does not affect binding to the other. This is a key distinction between the monospecific and bispecific antibodies – when the receptor levels are imbalanced, the monospecific antibodies demonstrate greater binding to the receptor in excess, while the bispecific antibody binds more of the limited receptor [**Figure 7C**]. However, the distinctions between monospecific antibody combinations and bispecific antibodies disappear at saturating antibody concentrations, where full receptor occupancy is achieved in both scenarios.

The distinction between monospecific antibody combinations and bispecific antibodies at sub-saturating concentrations is potentially very significant for therapeutic design in the context of heterogeneous expression of receptors. Tissues comprise a mix of cells expressing a single receptor, neither, or both (in various ratios). Individual cells also show significant heterogeneity in membrane receptor expression, particularly cancerous cells with mutations that alter gene expression. Both IL-6R and IL-8R are known to be overexpressed in cancer [12,28], and IL-6R has been quantified around 10^3^ receptors per cell in different carcinoma cell lines [65]. IL-8R has not been directly quantified in solid tumors, but it is found on monocytes and neutrophils at levels of 10^4^ to 10^5^ receptors per cell, and it was shown to be up-regulated relative to IL-6R in primary breast cancer tumor samples [28]. If an imbalance in receptor levels is present, it could be important to increase the apparent affinity for the less expressed receptor to achieve sufficient inhibition of the system. BS1 demonstrated stronger inhibition of migration and decreased metastatic burden *in vivo* compared to the combination of tocilizumab and 10H2 [28], and our results suggest that avidity effects may contribute to the underlying mechanism behind the superior performance of the bispecific antibody.

While our model provides significant insight into the mechanisms of bivalent and bispecific antibody binding, expanding the model to include additional processes and other cell types has the potential to give even more insight into these treatments. Physiologically, monoclonal antibodies are ultimately eliminated from the body via receptor-mediated endocytosis and subsequent intracellular catabolism [66], and higher affinity antibody binding leads to greater rates of endocytosis and degradation [67]. While receptor synthesis, internalization, and degradation were assumed to be negligible in modeling antibodies binding to cultured cells, given the importance of antibody-receptor complex internalization in determining the drug concentration profile and localization within tissues [67–69], it would be informative to extend the bivalent binding model presented here to include these processes. This modeling could in turn identify optimal bispecific design parameters to balance avidity with the rate of antibody endocytosis and elimination from the system. Additionally, as the aim of BS1 treatment is to inhibit IL-6/IL-8-driven metastasis, adding IL-6 and IL-8 secretion and binding to this system will allow us to directly model how antibody binding leads to ligand inhibition.

Our simulation results showed that receptor expression is critical for bispecific antibody binding, both in terms of total receptor levels and in the relative amounts of the individual targets. As IL-6 and IL-8 are both pleiotropic immune factors, their receptors are also expressed on healthy white blood cells and other tissues. Modeling multiple cell types with different receptor levels could allow us to quantify how the affinity and avidity effects presented here impact target tissue selectivity and further clarify differences between monospecific and bispecific antibodies [23,24]. Based on our current results, we hypothesize that cells that express a high concentration of only one receptor may act as “sinks” for monospecific antibodies, while the bispecific antibodies could preferentially bind to cells that express both receptors. Beyond local tissue binding, target expression on healthy cells could also impact antibody distribution and clearance from the body. Pharmacokinetic models of bispecific antibodies have previously been described [69,70], and extending our model to include circulation of the drug and its transport into the target tissue would enable study of the full treatment dynamics for monospecific versus bispecific antibodies.

## CONCLUSION

The binding of bispecific antibodies is governed by the intricate relationships between inherent binding affinity, combined multivalent avidity, therapeutic concentration, and target expression. Here we presented a mechanistic, computational model for antibodies targeting IL-6R and IL- 8R, comprised of a series of ordinary differential equations describing antibody binding dynamics. We fully parameterized the model from existing data, and our simulations closely match experimental data of monospecific and bispecific antibodies binding to cells expressing different levels of the IL-6 and IL-8 receptors. Model results describe the system dynamics and reveal key mechanisms underlying bispecific antibody behavior. The model also predicts the consequent receptor occupancy due to the antibodies, as well as the distribution of receptors into binary (antibody-receptor) and ternary (receptor-antibody-receptor) complexes; ultimately, the impact on receptor complex formation, rather than the amount of receptor binding, is critical for therapeutic performance. We observed that the bispecific antibody studied demonstrates strong cross-linking and avidity effects, which increase receptor residence time. Ternary complex formation is maximized when binding affinity is balanced with antibody concentration (for both monospecific and bispecific antibodies) and target expression level. When the target receptors are present in unequal amounts, monospecific and bispecific antibodies demonstrate distinct binding patterns – monospecific antibodies bind more strongly to the excess target, whereas bispecific antibodies show greater apparent affinity for the limited target at sub-saturating concentrations. Overall, our quantitative model of anti-IL-6R/anti-IL-8R antibodies provides clear mechanistic insight into the dynamics of homo- and heterobivalent antibodies and leads to actionable predictions of optimal therapeutic design for maximal binding. The results provided here include specific parameter values for these antibodies for IL-6R and IL-8R, but many of the insights can be applied generally to other bispecific antibodies, and the model itself can be repurposed to analyze other therapeutic systems of interest.

## DATA AND CODE AVAILABILITY

All code written in support of this publication is publicly available at https://github.com/christyray/bispecific-binding-model, and we have archived our code on Zenodo (DOI: <u>10.5281/zenodo.10396436</u>). Experimental data, simulation input files, and generated data are available on Zenodo at https://doi.org/10.5281/zenodo.10393562. Code for model simulations is written in MATLAB using version R2022a.

