## Supplemental Information for "Mechanistic computational modeling of monospecific and bispecific antibodies targeting interleukin-6/8 receptors"

### *Binding Model Equations*

α is a unit conversion parameter, used in Equations S2, S5, S8 to convert the rate terms from (# receptors per cell)$^{-1}$ s$^{-1}$ to nM$^{-1}$ s$^{-1}$. It is defined in Equation S1 where $n_{cell}$ is the number of cells in the experiment or simulation, V is the volume containing the cells and the antibodies, and $N_A$ is Avogadro's number, 6.022 × $10^{23}$ molecules per mole. For the conditions used in the flow cytometry binding assays and in the model simulations, $n_{cell}$ = $10^5$ cells and V = 200 μL, making α = 8.3 × $10^{-7}$ nM/(# / cell).

$$\alpha = \frac{n_{cell}}{V * N_A} \qquad (S1)$$

$$\frac{d[Toci]}{dt} = \alpha * k_{off,Toci-6R}[Toci \cdot 6R] - 2 * \alpha * k_{on,Toci-6R}[Toci][6R] \qquad (S2)$$

$$\frac{d[Toci \cdot 6R]}{dt} = 2 * k_{on,Toci-6R}[Toci][6R] + 2 * k_{off,Toci-6R^*}[6R \cdot Toci \cdot 6R] - k_{on,Toci-6R^*}[6R][Toci \cdot 6R] - k_{off,Toci-6R}[Toci \cdot 6R] \qquad (S3)$$

$$\frac{d[6R \cdot Toci \cdot 6R]}{dt} = k_{on,Toci-6R^*}[6R][Toci \cdot 6R] - 2 * k_{off,Toci-6R^*}[6R \cdot Toci \cdot 6R] \qquad (S4)$$

$$\frac{d[H2]}{dt} = \alpha * k_{off,H2-8R}[H2 \cdot 8R] - 2 * \alpha * k_{on,H2-8R}[H2][8R] \qquad (S5)$$

$$\frac{d[H2 \cdot 8R]}{dt} = 2 * k_{on,H2-8R}[H2][8R] + 2 * k_{off,H2-8R^*}[8R \cdot H2 \cdot 8R] - k_{on,H2-8R^*}[8R][H2 \cdot 8R] - k_{off,H2-8R}[H2 \cdot 8R] \qquad (S6)$$

$$\frac{d[8R \cdot H2 \cdot 8R]}{dt} = k_{on,H2-8R^*}[8R][H2 \cdot 8R] - 2 * k_{off,H2-8R^*}[8R \cdot H2 \cdot 8R] \quad (S7)$$

$$\frac{d[BS1]}{dt} = \alpha * k_{off,BS1-6R}[BS1 \cdot 6R] + \alpha * k_{off,BS1-8R}[BS1 \cdot 8R] - \alpha * k_{on,BS1-6R}[BS1][6R] - \alpha * k_{on,BS1-8R}[BS1][8R] \quad (S8)$$

$$\frac{d[BS1 \cdot 6R]}{dt} = k_{on,BS1-6R}[BS1][6R] + k_{off,BS1-8R^*}[6R \cdot BS1 \cdot 8R] - k_{on,BS1-8R^*}[8R][BS1 \cdot 6R] - k_{off,BS1-6R}[BS1 \cdot 6R] \quad (S9)$$

$$\frac{d[BS1 \cdot 8R]}{dt} = k_{on,BS1-8R}[BS1][8R] + k_{off,BS1-6R^*}[6R \cdot BS1 \cdot 8R] - k_{on,BS1-6R^*}[6R][BS1 \cdot 8R] - k_{off,BS1-8R}[BS1 \cdot 8R] \quad (S10)$$

$$\frac{d[6R \cdot BS1 \cdot 8R]}{dt} = k_{on,BS1-6R^*}[6R][BS1 \cdot 8R] + k_{on,BS1-8R^*}[8R][BS1 \cdot 6R] - k_{off,BS1-6R^*}[6R \cdot BS1 \cdot 8R] - k_{off,BS1-8R^*}[6R \cdot BS1 \cdot 8R] \quad (S11)$$

$$\frac{d[6R]}{dt} = k_{off,Toci-6R}[Toci \cdot 6R] + 2 * k_{off,Toci-6R^*}[6R \cdot Toci \cdot 6R] + k_{off,BS1-6R}[BS1 \cdot 6R] + k_{off,BS1-6R^*}[6R \cdot BS1 \cdot 8R] - 2 * k_{on,Toci-6R}[Toci][6R] - k_{on,Toci-6R^*}[6R][Toci \cdot 6R] - k_{on,BS1-6R}[BS1][6R] - k_{on,BS1-6R^*}[6R][BS1 \cdot 8R] \quad (S12)$$

$$\frac{d[8R]}{dt} = k_{off,H2-8R}[H2 \cdot 8R] + 2 * k_{off,H2-8R^*}[8R \cdot H2 \cdot 8R] + k_{off,BS1-8R}[BS1 \cdot 8R] + k_{off,BS1-8R^*}[6R \cdot BS1 \cdot 8R] - 2 * k_{on,H2-8R}[H2][8R] - k_{on,H2-8R^*}[8R][H2 \cdot 8R] - k_{on,BS1-8R}[BS1][8R] - k_{on,BS1-8R^*}[8R][BS1 \cdot 6R] \quad (S13)$$

### *Normalization-Schemes*

Optimization of the binding model rate constants with the different normalization schemes (detailed in **Table 3**) yields similar optimized parameter sets, with slightly different parameter value distributions for each scheme [**Figure S4**]. For the optimizations normalized to the BS1 data at binding saturation in particular, the optimizations converged to a single parameter set at a very high frequency [**Figure S4A**].

While the optimizations with normalization against the individual antibodies do still contain a most frequent optimal point, they also yield a wider variety of possible parameter values [**Figure S4B**]. Each of the distributions shows two or more commonly reoccurring values, and the lowest cost parameter value is not always the same as the most frequent value. Some of the parameter values are similar to the values from the normalization against BS1, but the $k_{on,6R^*}$ and the $k_{off,8R}$ values differ by almost two orders of magnitude.

Further, optimizations performed with normalization against the BS1 data, both normalized to the bound concentrations at binding saturation and normalized to the maximum bound concentration, demonstrate a greater proportion of low-cost optimal sets than the optimizations normalized to each antibody separately, as illustrated by the cumulative distribution of the cost of each set of optimal values [**Figure S5**]. The greater variance in optimal values for the normalization to individual antibodies separately is likely due to a reduction in data meaningfulness when comparisons between antibodies are lost. Many of the parameter sets from the different normalization schemes match well to the experimental data, but a few sets show more substantial variation [**Figure S7**].

In total, these results demonstrate that ideal normalization scheme for the model output is normalization to the amount of bound BS1 at binding saturation (i.e., “BS1, Data”). This scheme showed a strong convergence to a single optimal parameter set, and simulations with this optimal parameter set replicate the experimental data well. For these reasons, this normalization scheme was selected as the primary normalization method for the model analysis as described in the *Results*.

***Table S1.*** *Original model parameters and their relationship to the simplified parameters after applying the model assumptions.*

| Original | Simplified | Best Fit Value | Units |
|---|---|---|---|
| $k_{on,Toci-6R}$ | $k_{on,6R}$ | $5.92 \times 10^{-6}$ | $nM^{-1}s^{-1}$ |
| $k_{on,Toci-6R^*}$ | $k_{on,6R^*}$ | $8.11 \times 10^{-8}$ | $\left(\frac{\#}{cell}\right)^{-1} s^{-1}$ |
| $k_{off,Toci-6R}$ | $k_{off,6R}$ | $5.61 \times 10^{-5}$ | $s^{-1}$ |
| $k_{off,Toci-6R^*}$ | $k_{off,6R}$ | $5.61 \times 10^{-5}$ | $s^{-1}$ |
| $k_{on,H2-8R}$ | $k_{on,8R}$ | $9.03 \times 10^{-6}$ | $nM^{-1}s^{-1}$ |
| $k_{on,H2-8R^*}$ | $k_{on,8R^*}$ | $1.24 \times 10^{-7}$ | $\left(\frac{\#}{cell}\right)^{-1} s^{-1}$ |
| $k_{off,H2-8R}$ | $k_{off,8R}$ | $6.38 \times 10^{-5}$ | $s^{-1}$ |
| $k_{off,H2-8R^*}$ | $k_{off,8R}$ | $6.38 \times 10^{-5}$ | $s^{-1}$ |
| $k_{on,BS1-6R}$ | $k_{on,6R}$ | $5.92 \times 10^{-6}$ | $nM^{-1}s^{-1}$ |
| $k_{on,BS1-8R}$ | $k_{on,8R}$ | $9.03 \times 10^{-6}$ | $nM^{-1}s^{-1}$ |
| $k_{on,BS1-6R^*}$ | $k_{on,6R^*}$ | $8.11 \times 10^{-8}$ | $\left(\frac{\#}{cell}\right)^{-1} s^{-1}$ |
| $k_{on,BS1-8R^*}$ | $k_{on,8R^*}$ | $1.24 \times 10^{-7}$ | $\left(\frac{\#}{cell}\right)^{-1} s^{-1}$ |
| $k_{off,BS1-6R}$ | $k_{off,6R}$ | $5.61 \times 10^{-5}$ | $s^{-1}$ |
| $k_{off,BS1-8R}$ | $k_{off,8R}$ | $6.38 \times 10^{-5}$ | $s^{-1}$ |

| Original | Simplified | Best Fit Value | Units |
|---|---|---|---|
| $k_{off,BS1-6R^*}$ | $k_{off,6R}$ | $5.61 \times 10^{-5}$ | $s^{-1}$ |
| $k_{off,BS1-8R^*}$ | $k_{off,8R}$ | $6.38 \times 10^{-5}$ | $s^{-1}$ |

***Table S2.*** *IL-6R and IL-8R binding affinities for the monospecific and bispecific antibodies calculated directly from in vitro HEK 293T cell surface binding assays. Values are given as dissociation constants ($K_D$) in nM and were originally reported by Yang et al [28].*

| | **IL-6R$^+$ / IL-8R$^-$** | **IL-6R$^-$ / IL-8R$^+$** | **IL-6R$^+$ / IL-8R$^+$** |
|---|---|---|---|
| Tocilizumab | 3.1 | N/A | 3.1 |
| 10H2 | N/A | 3.6 | 4.1 |
| BS1 | 24 | 10 | 14 |

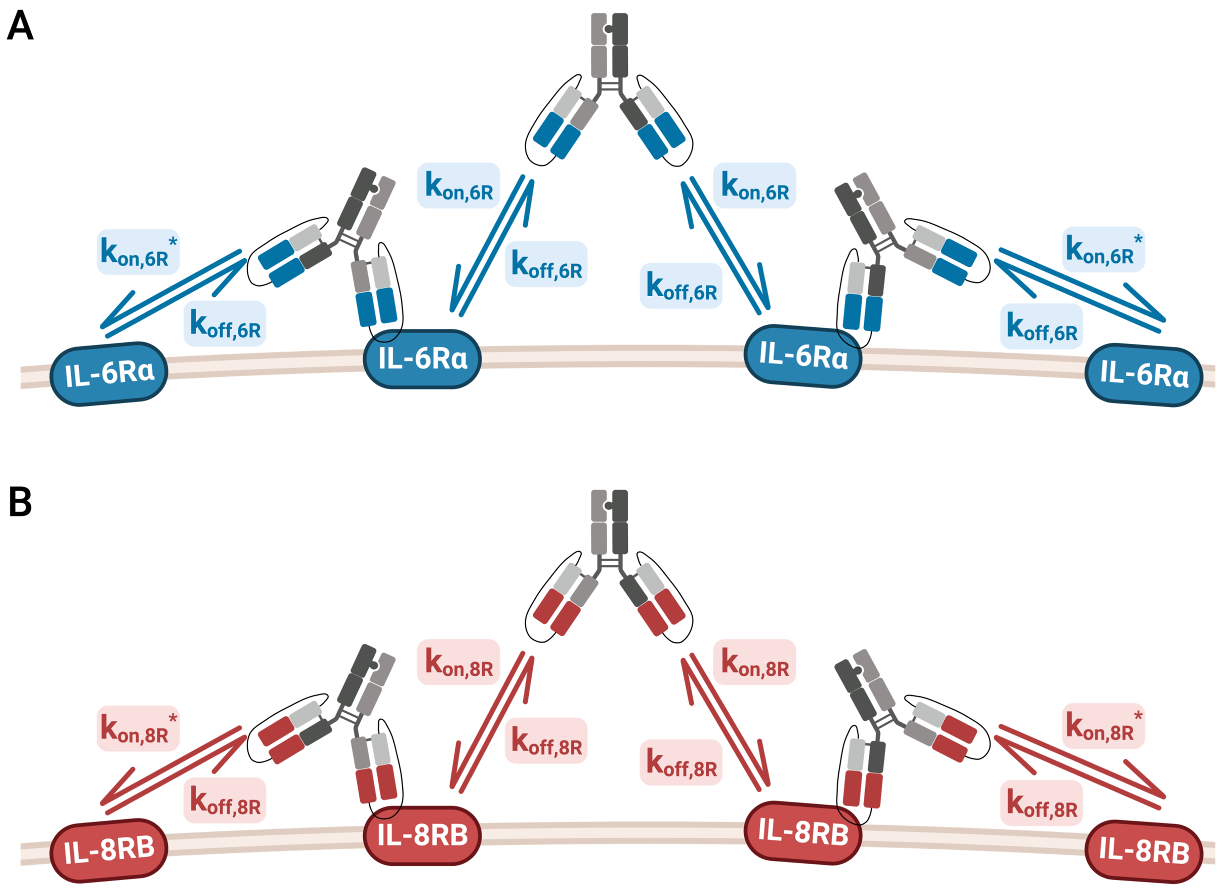


**Figure S1. Monoclonal antibody binding model kinetics.** Schematic of the IL-6Rα/IL-8RB antibody-binding model for the two monoclonal antibodies, tocilizumab (anti-IL-6Rα) (**A**) and 10H2 (anti-IL-8RB) (**B**). As in the BS1 binding model [**Figure 1B**], $k_{on,6R}$ and $k_{on,8R}$ describe the association rates for the formation of binary antibody-receptor complexes, and $k_{on,6R^*}$ and $k_{on,8R^*}$ describe the association rates for the formation ternary receptor-antibody-receptor complexes. The same $k_{off,6R}$ and $k_{off,8R}$ rate constants are used for the dissociation of both the binary and the ternary complexes. Notably, in this study we simplify the parameter optimization by assuming that the binding rate constants for the bispecific antibody [**Figure 1B**] are the same as the equivalent reactions for the monospecific antibodies. This figure was created with BioRender.com.

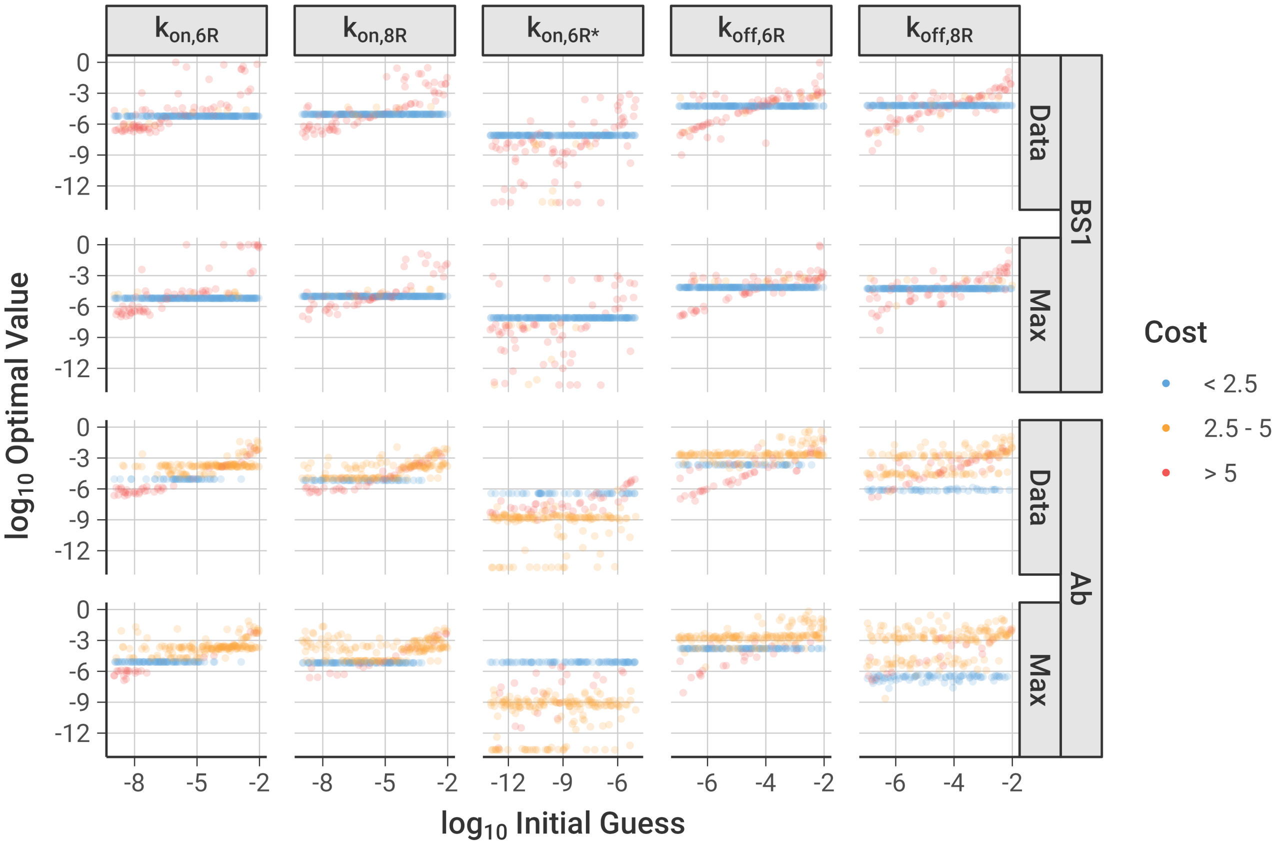


**Figure S2. Relationship between initial guesses and optimized values for each binding reaction rate constant, separated by normalization options used.** ***BS1*** describes simulations that were normalized against the concentration of bound BS1 at the end time point, and ***Ab*** indicates simulations that were normalized against the concentration of that specific antibody at the end time point. ***Data*** depicts simulations that were normalized using the bound concentrations at the binding saturation, as was done for the experimental data, and ***Max*** describes simulations that were normalized using the bound concentration at the maximum initial antibody concentration.

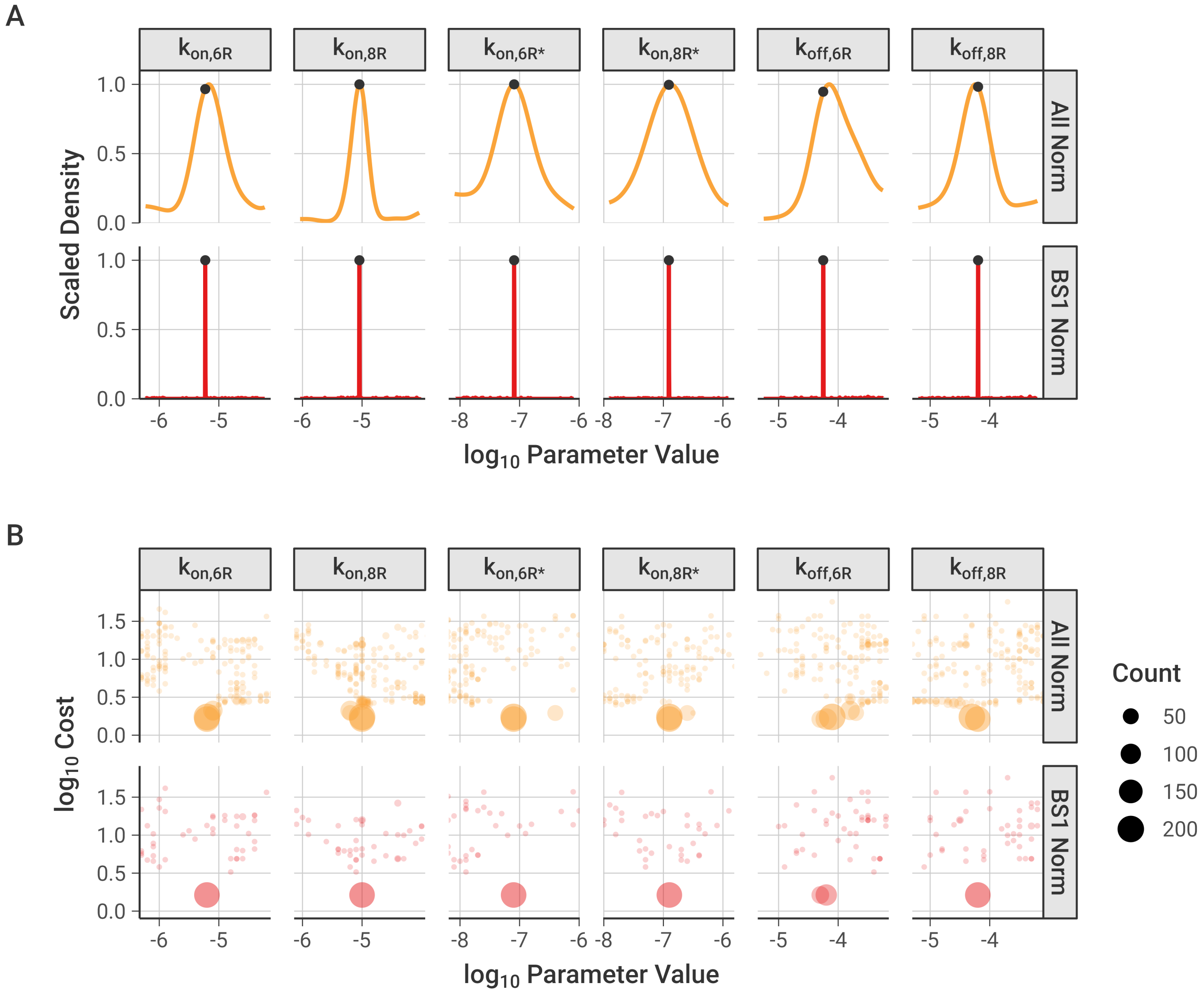


**Figure S3. Frequency and cost of optimized binding model parameter sets, showing a limited range of values around the lowest-cost parameter set.** To better visualize the distribution of the parameter sets, the plotted values are limited to one order of magnitude above and below the values from the lowest cost parameter set [**Table 2**]. **A,** Distribution of optimized parameter values across all optimizations performed, with marked points indicating the values of the lowest cost parameter set. **B,** Relationship between optimized parameter values and the cost of the optimized parameter sets compared to experimental data, separated by parameter. Optimized points with the same value are grouped into a single point, with the point size indicating how many optimized parameter values are in the group.

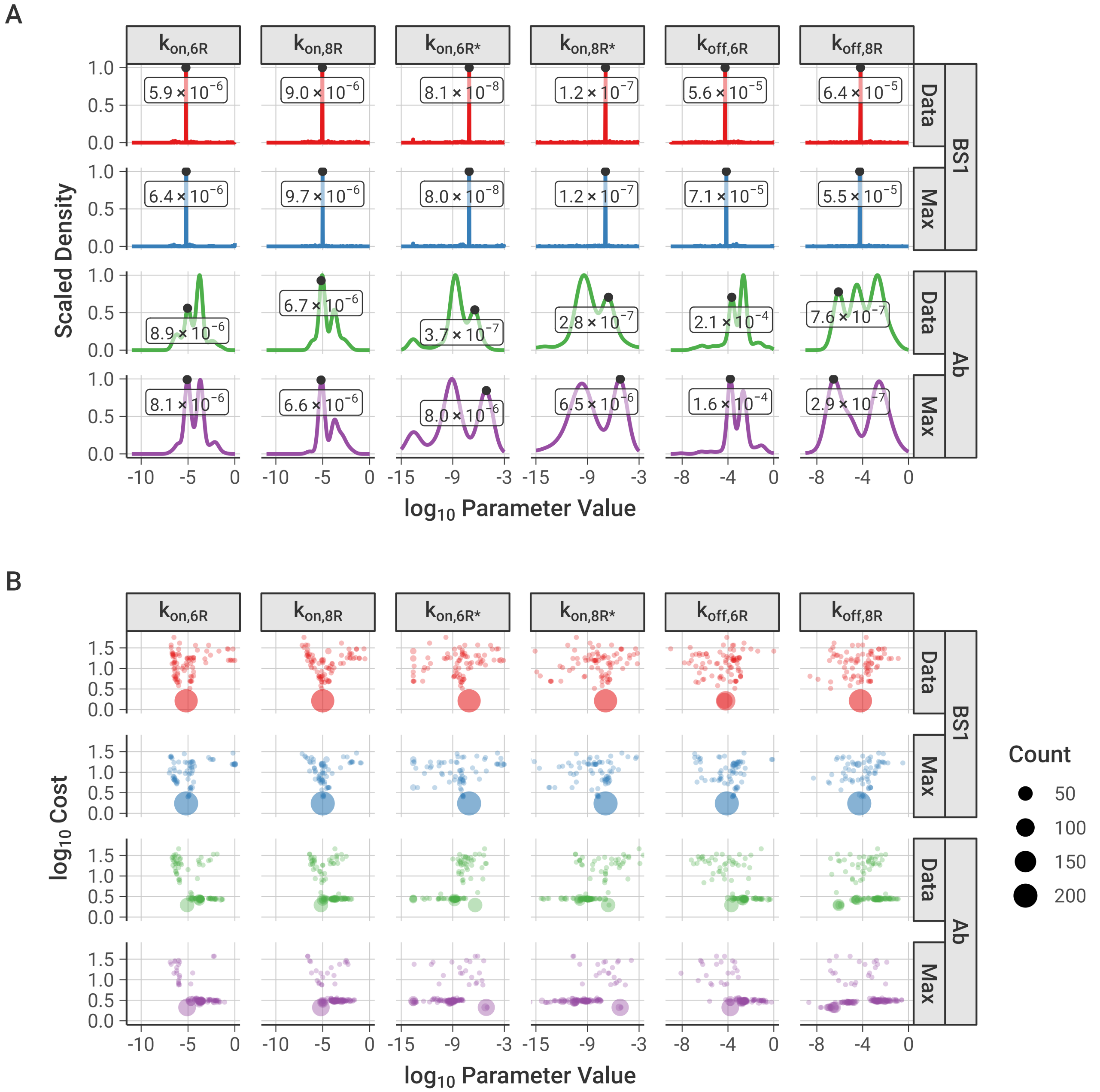


**Figure S4. Distribution of optimized parameter values and cost compared to experimental data, separated by parameter and normalization type.** Normalization types are separated in the same way as for the initial guesses [**Figure S2**]. **A,** Distribution of optimized parameter values across all optimizations performed, separated by normalization options. Marked points and corresponding labels indicate the values of the lowest cost parameter set for those specific normalization options. **B,** Relationship between optimized parameter values and cost compared to experimental data, separated by parameter. Optimized points with the same value are grouped into a single point, with the point size indicating how many optimized parameters are in the group.

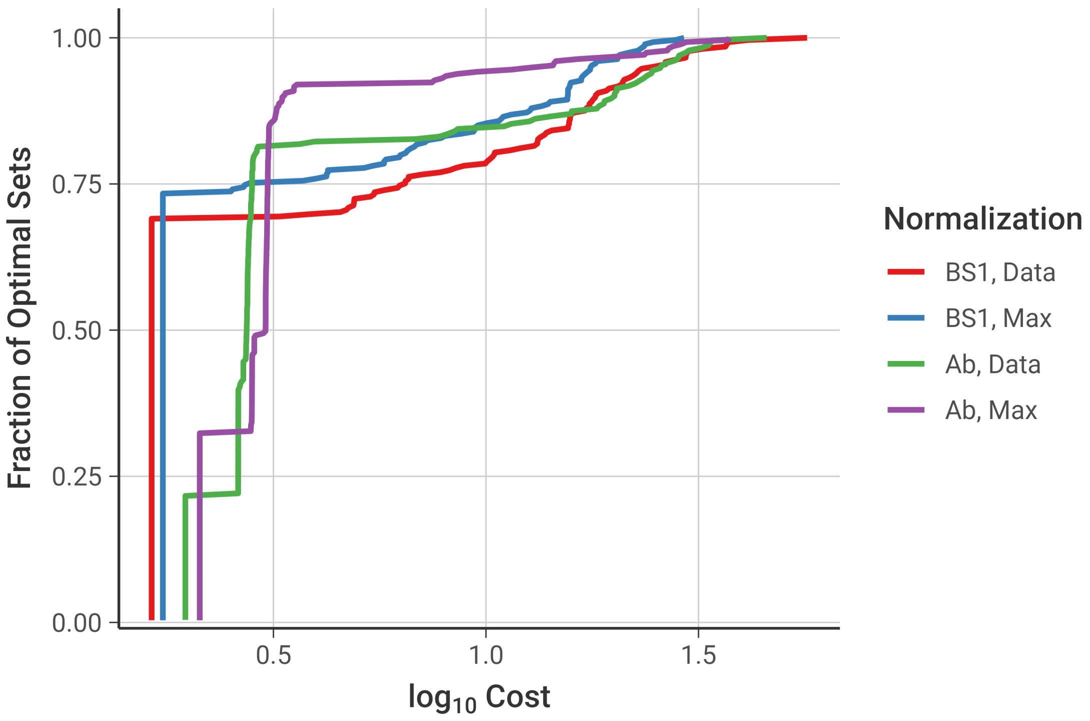


**Figure S5.** Cumulative distribution of the cost of the optimized parameter sets, separated by normalization options used. The curves depict the fraction of optimal parameter sets that were below a given cost value. Parameter sets where the optimization did not converge were omitted.

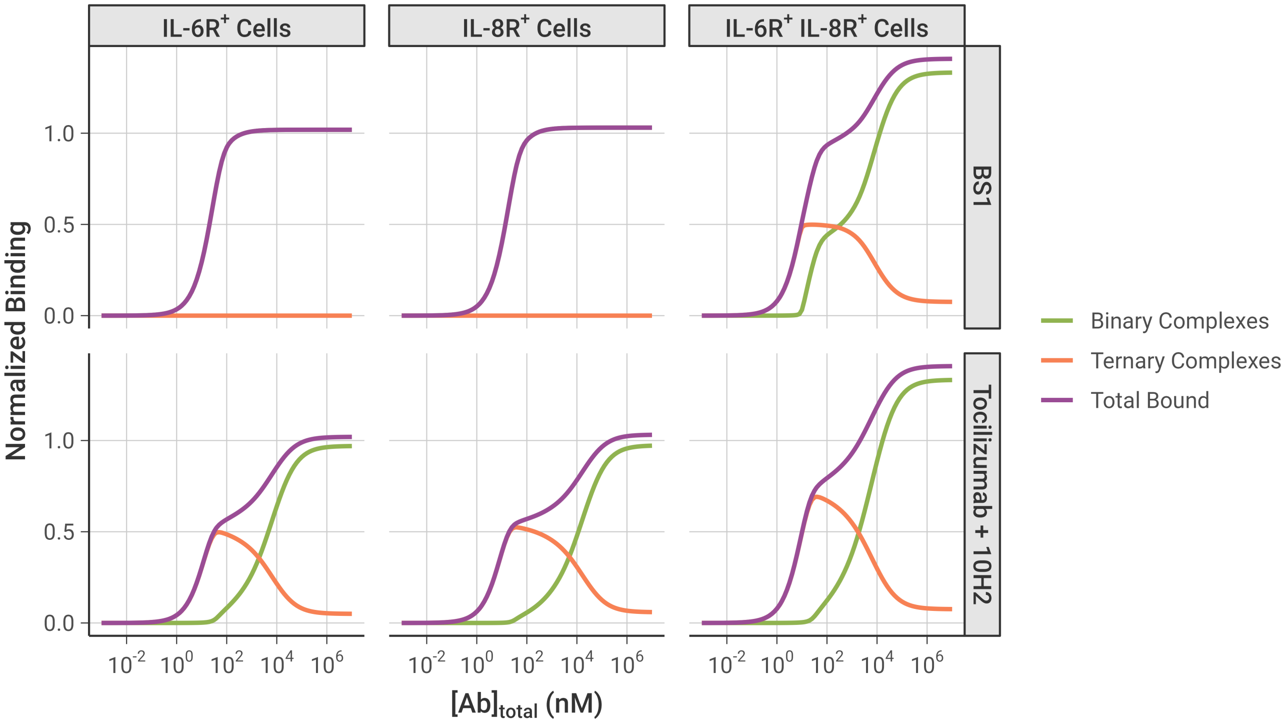


**Figure S6. Comparison of normalized binding curves between the different antibodies**. Bound concentrations are divided among binary complexes, ternary complexes, and total bound antibody. Simulations were performed under the same conditions as the binding experiments: $10^5$ cells/well, receptor expression levels from the transduced cell lines [**Table 1**], and with a 2-hour initial association period followed by a 15-minute free antibody washout. Model output is normalized to the bound concentration of BS1 at the same initial antibody concentrations used to normalize the experimental data.

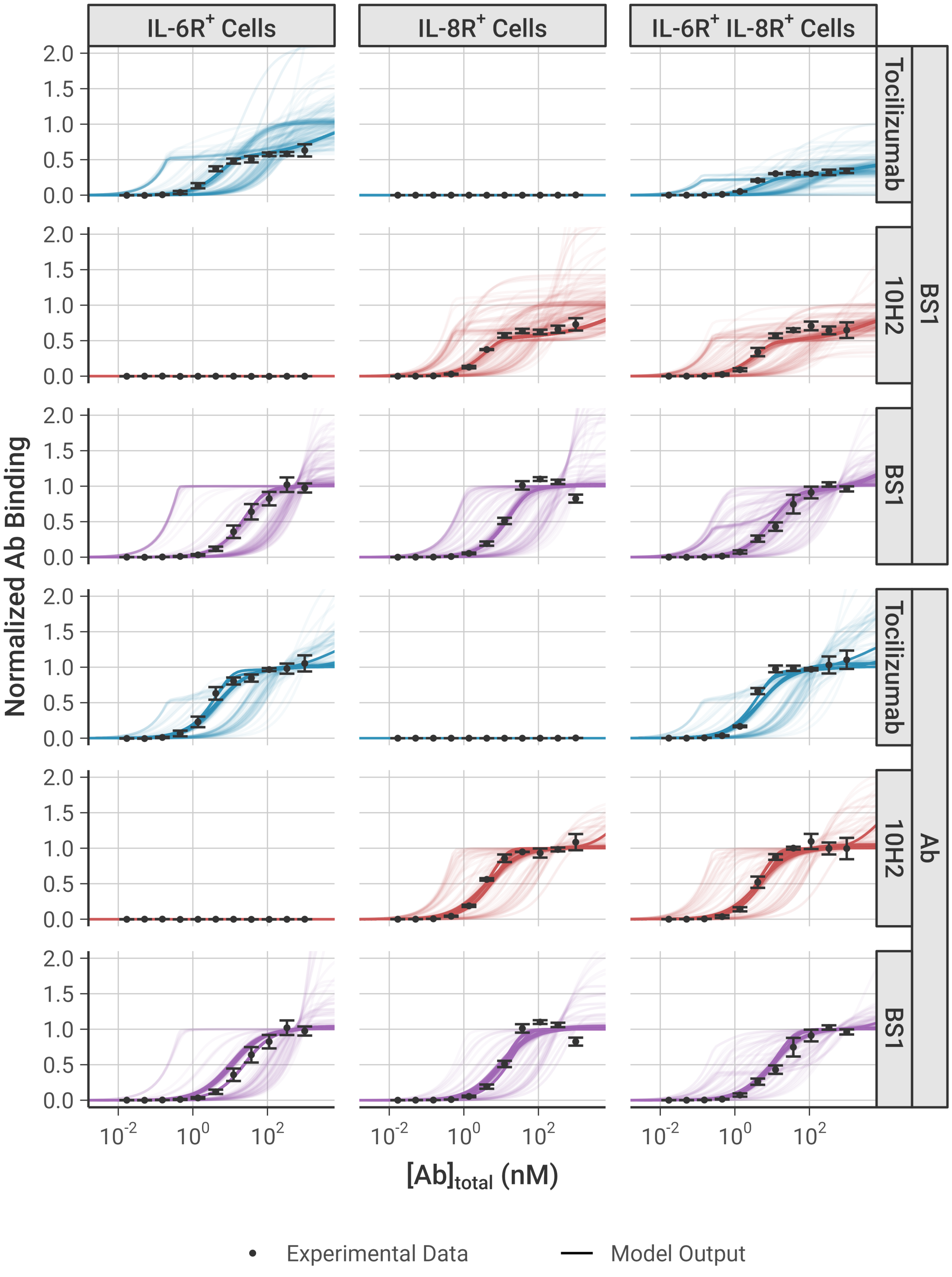


**Figure S7. Model simulation results using each of the optimized parameter sets compared to the experimental data used to fit the model parameters.** Simulations were performed under the same conditions as the experiment, with a 2-hour initial binding period followed by a 15-minute antibody washout and with the experimental receptor expression levels [**Table 1**]. The model simulation results (lines) are compared to the equivalent experimental data (points), and each optimized parameter set is represented with a separate line. Panels are separated by the normalization basis used, with “BS1” referring to simulations and experimental data normalized against the bound BS1 concentration in all cell lines and “Ab” referring simulations and experimental data where each antibody was normalized against itself. Each panel depicts all optimal parameter sets obtained with that particular normalization method. Model output and experimental data are each normalized to the output/data from the concentrations where binding reached saturation. The error bars depict the standard error from three experimental replicates.

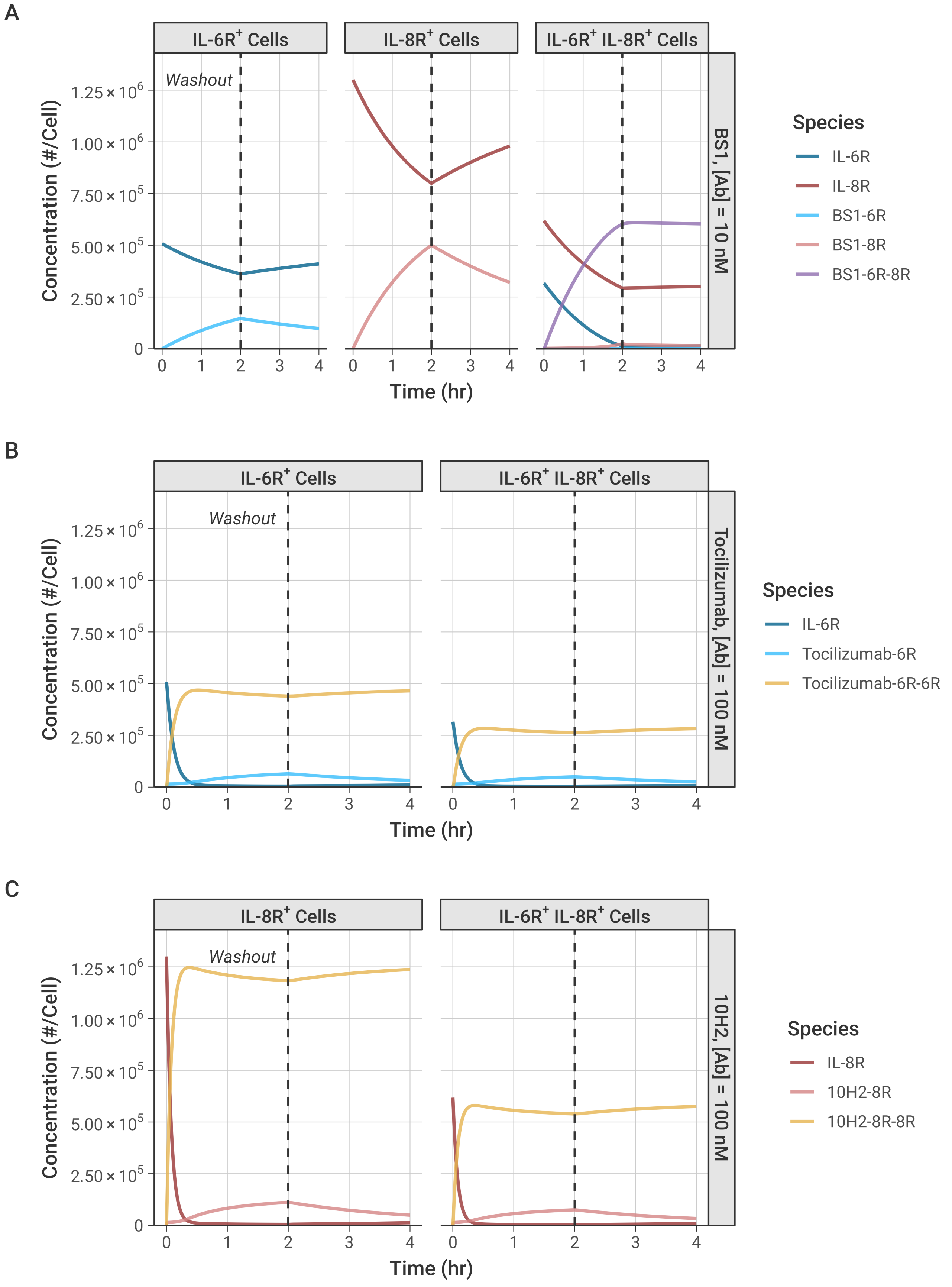
A
IL-6R+ Cells
IL-8R+ Cells
IL-6R+ IL-8R+ Cells
Washout
BS1, [Ab] = 10 nM
Concentration (#/Cell)
Time (hr)
Species
IL-6R
IL-8R
BS1-6R
BS1-8R
BS1-6R-8R
B
IL-6R+ Cells
IL-6R+ IL-8R+ Cells
Washout
Tocilizumab, [Ab] = 100 nM
Concentration (#/Cell)
Time (hr)
Species
IL-6R
Tocilizumab-6R
Tocilizumab-6R-6R
C
IL-8R+ Cells
IL-6R+ IL-8R+ Cells
Washout
10H2, [Ab] = 100 nM
Concentration (#/Cell)
Time (hr)
Species
IL-8R
10H2-8R
10H2-8R-8R

**Figure S8. Simulations quantify bivalent antibody binding to IL-6R and IL-8R over time.** Cell number = $1 \times 10^5$ for all simulations. Free antibody concentration was set to 0 nM at 2 hours to simulate antibody washout from the system. The expressions of IL-6R and IL-8R from the transduced experimental cell lines were used in the simulations [**Table 1**]. **A,** Simulations with an initial BS1 concentration of 10 nM, compared to the simulations with 100 nM of BS1 shown in the main text [**Figure 4**]. **B,** Simulations of tocilizumab at an initial concentration of 100 nM in the IL-6R$^+$ cell lines. **C,** Simulations of 10H2 at an initial concentration of 100 nM in the IL-8R$^+$ cell lines.

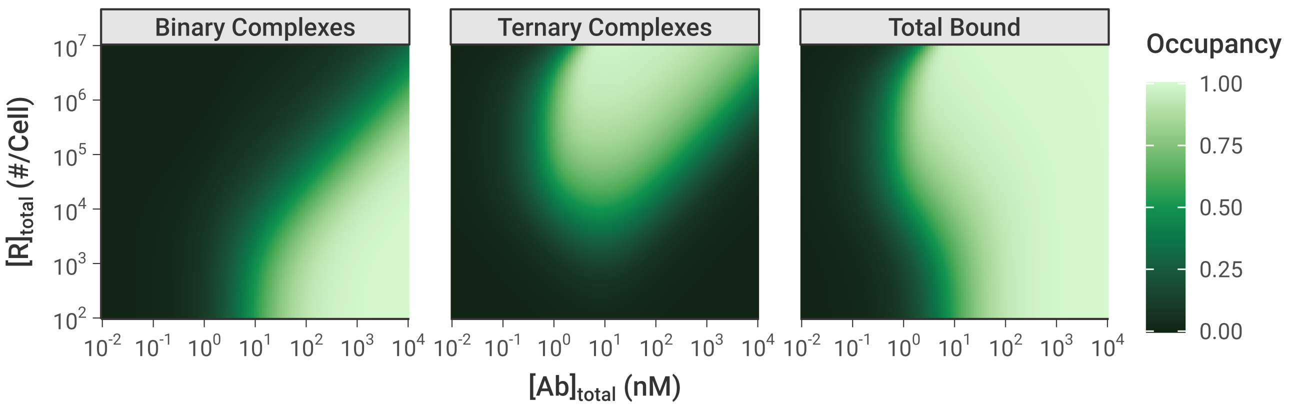


**Figure S9. Simulated Binary (Ab-R), Ternary (R-Ab-R), and Total Bound (Binary + Ternary) concentrations of Antibody-Receptor complexes using the combination of tocilizumab and 10H2.** IL-6R and IL-8R are present in a 1:1 ratio, as are tocilizumab and 10H2. Simulations were performed for 24 hours after antibody dosing. The color indicates the fraction of the total receptor (IL-6R + IL-8R) that is bound in each antibody-receptor complex type.

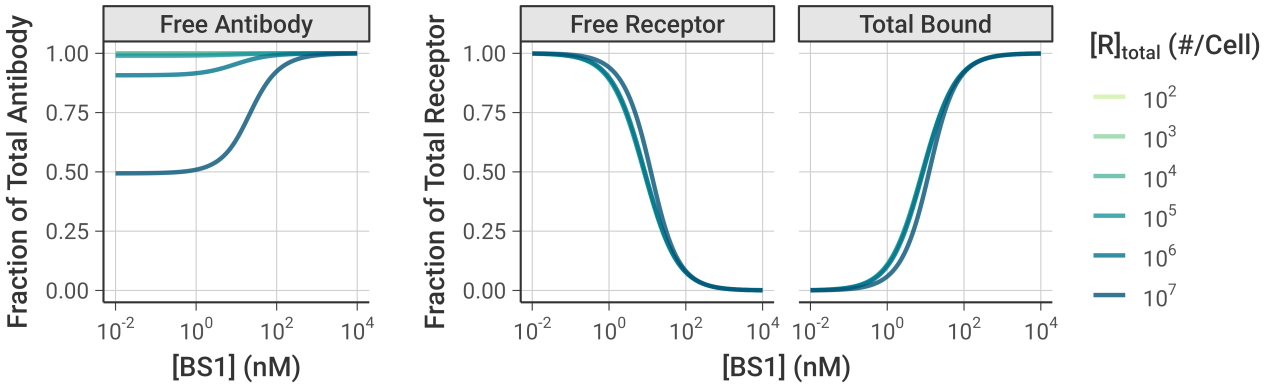


**Figure S10. Simulations of monovalent BS1 binding.** Fraction of total BS1 and receptor concentrations free and bound over varying initial BS1 concentration. The first panel shows the fraction of total BS1 concentration that is unbound, and the other panels show the fraction of total receptor concentration (IL-6R + IL-8R) that is unbound and bound. The association rate constants for the formation of ternary complexes ($k_{on,6R^*}$ and $k_{on,8R^*}$) were set to 0 to restrict BS1 to monovalent binding only. IL-6R and IL-8R are present in a 1:1 ratio, and simulations were performed for 24 hours after antibody dosing.

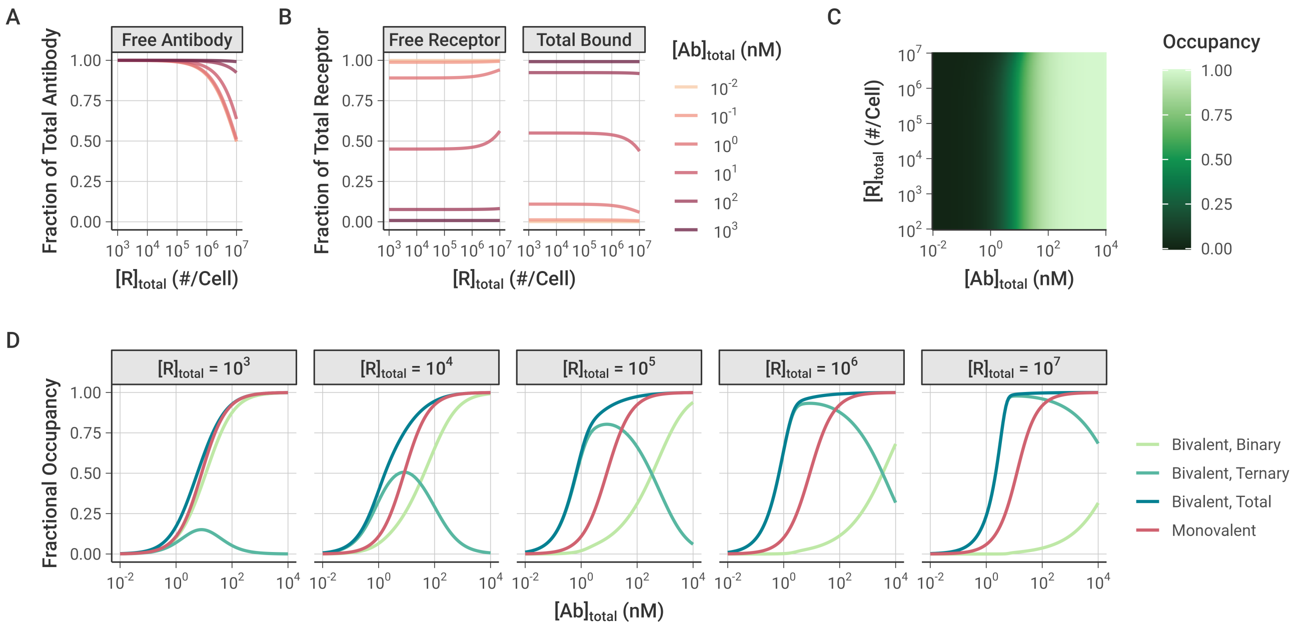


**Figure S11. Simulations of the combination of tocilizumab and 10H2 restricted to monovalent binding only.** Simulations were performed for over varying initial antibody and receptor concentrations. The association rate constants for the formation of ternary complexes ($k_{on,6R^*}$ and $k_{on,8R^*}$) were set to 0 to restrict both antibodies to monovalent binding only. IL-6R and IL-8R are present in a 1:1 ratio, as were tocilizumab and 10H2, and simulations were performed for 24 hours after antibody dosing. The simulation conditions are the same as those shown for BS1 in the main text [**Figure 6**]. **A,** Fraction of total antibody concentration (tocilizumab + 10H2) that is free (unbound) for different levels of receptor expression and initial antibody concentration. **B,** Fraction of total receptor concentration (IL-6R + IL-8R) that is unbound (free) or bound (in binary antibody-receptor complexes) for different levels of receptor expression and initial total antibody (tocilizumab + 10H2) concentration. **C,** Heat map of bound receptor fraction over varying antibody and receptor concentrations. The color indicates the fraction of the total receptor (IL-6R + IL-8R) that is bound to antibody. **D,** Comparison of monovalent and bivalent binding. The lines indicate the fraction of total receptor (IL-6R + IL-8R) that is bound in different complex types in the original simulations and the simulations restricted to monovalent binding only. The panels are divided by the total receptor concentration (in # receptors/cell).

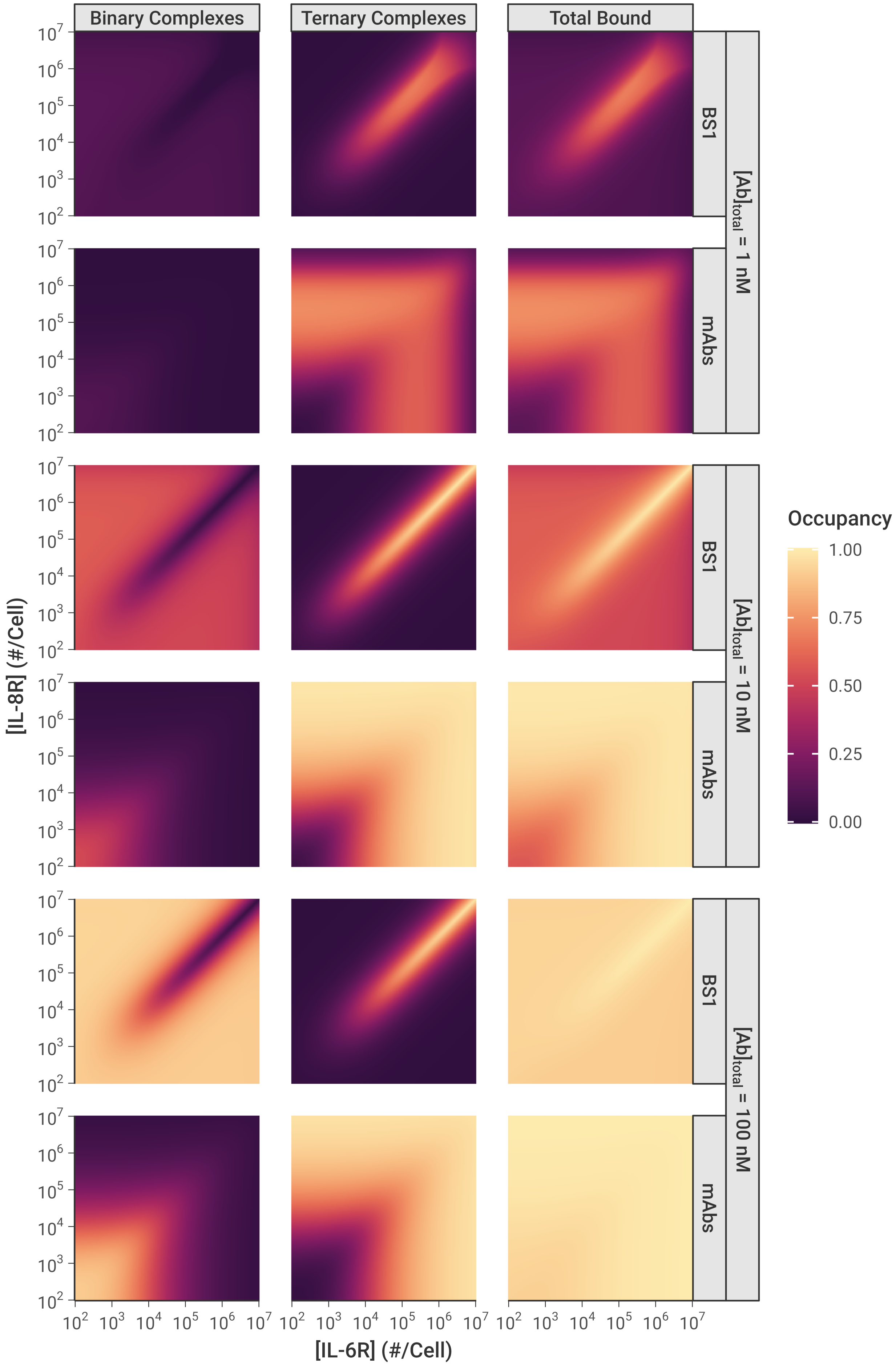

Binary Complexes
Ternary Complexes
Total Bound
BS1
mAbs
[Ab]total = 1 nM
[Ab]total = 10 nM
[Ab]total = 100 nM
Occupancy
1.00
0.75
0.50
0.25
0.00
[IL-8R] (#/Cell)
[IL-6R] (#/Cell)

**Figure S12. Fraction of receptors bound in Binary and Ternary complexes and Total Bound receptor (Binary + Ternary) across different IL-6R and IL-8R expression levels and initial antibody concentrations.** The color indicates the fraction of total receptor (IL-6R + IL-8R) that is bound in each antibody-receptor complex type. The antibody concentration is the total initial concentration of antibody in the system; “mAbs” refers to tocilizumab and 10H2 together in a 1:1 concentration ratio. Simulations were performed for 24 hours after antibody dosing. The panels for [Ab] = 10 nM were presented in the main text [**Figure 7A**] and are repeated here for comparison to the other concentrations.

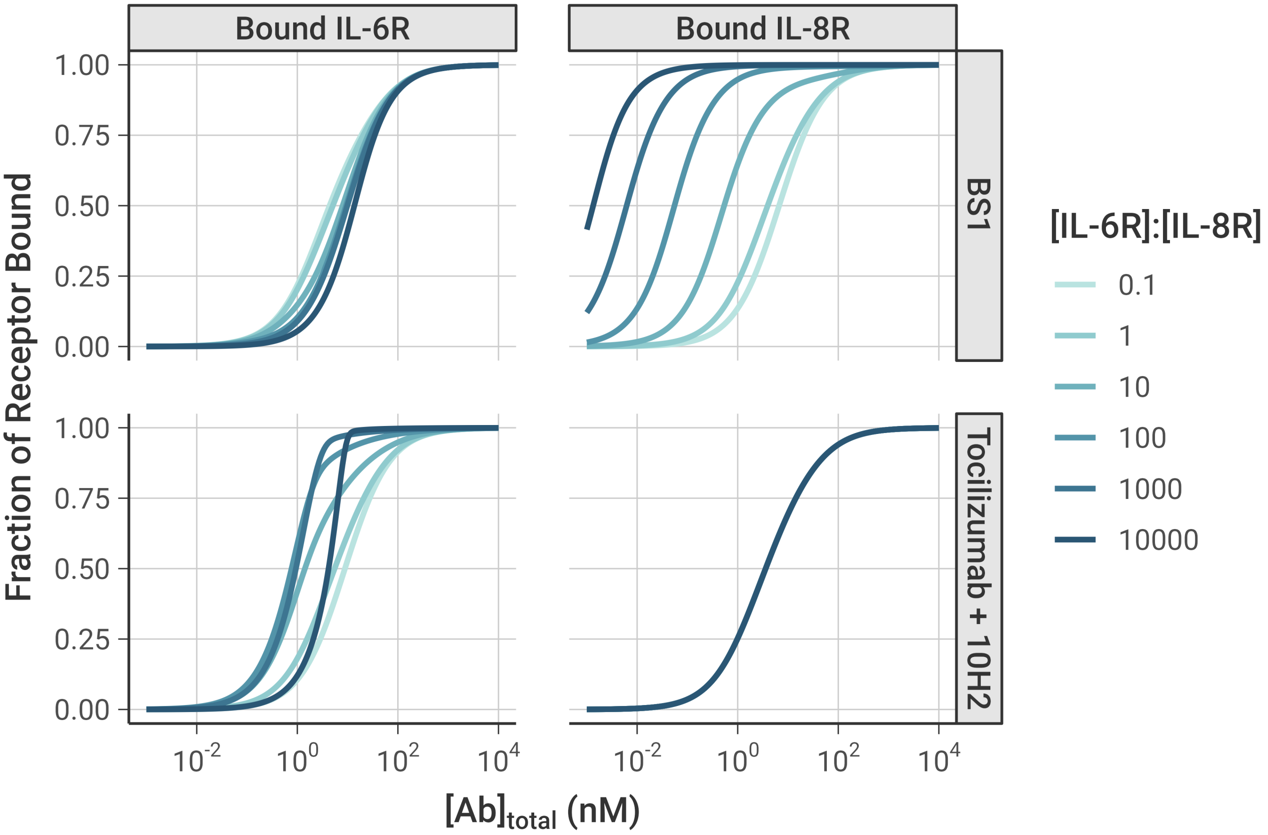


**Figure S13.** The fractional occupancy of each receptor individually when one receptor (IL-6R) is in excess. IL-8R was fixed at $10^3$ receptors/cell for these simulations, while IL-6R ranged from $10^2$ to $10^7$ receptors/cell. The fractional occupancy indicates the fraction of the specific receptor concentration (either IL-6R or IL-8R) that is bound to antibody (either BS1 or the combination of tocilizumab and 10H2). The fractional occupancy when IL-6R was fixed and IL-8R was in excess was shown in the main text [**Figure 7**]

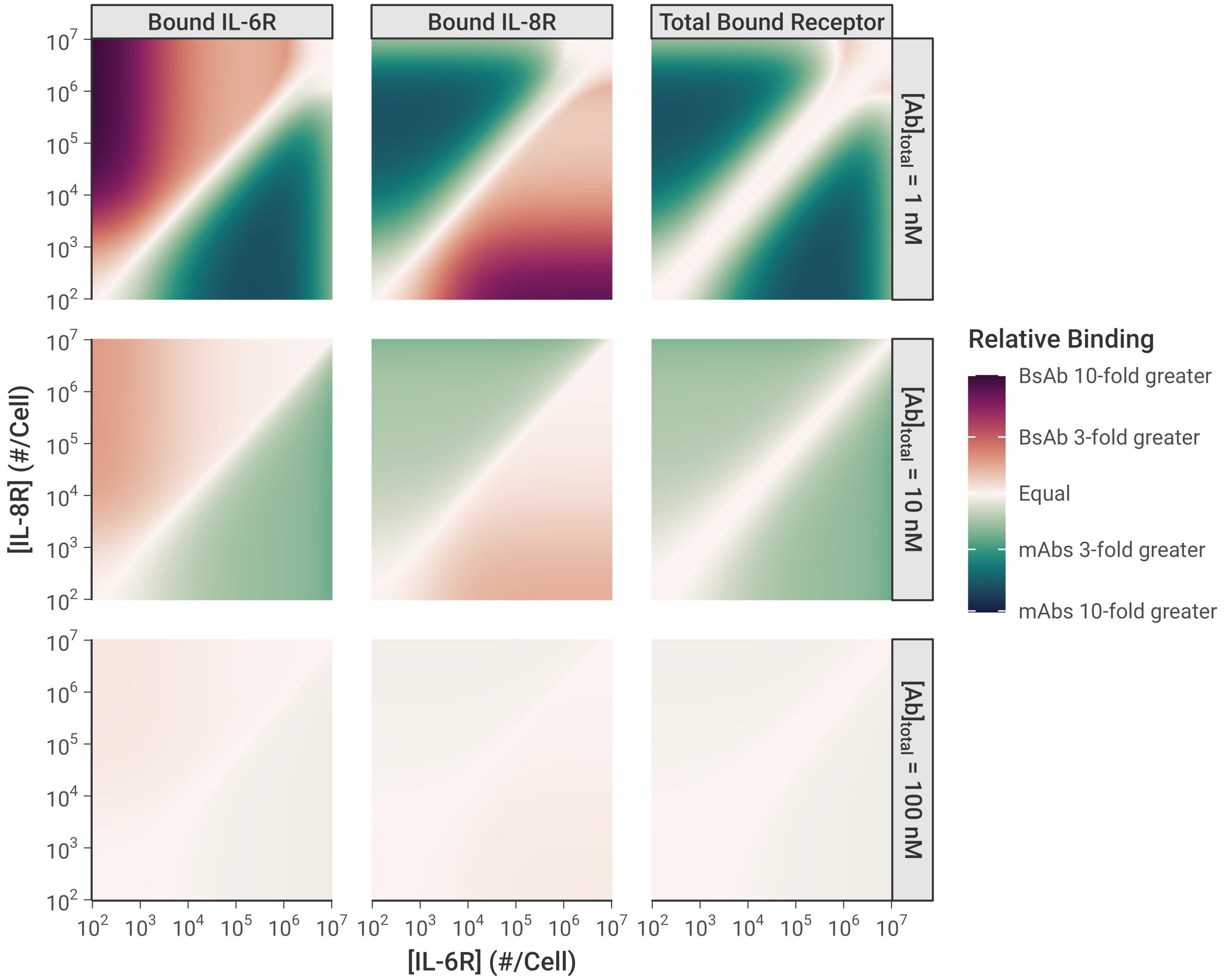


**Figure S14. Relative binding of BS1 and the combination of monoclonal antibodies across different IL-6R and IL-8R expression levels and initial total antibody concentrations.** The color indicates the relative bound receptor, where relative binding is the ratio of fractional bound receptor (the fraction of total IL-6R + IL-8R bound to antibody) when BS1 is used compared to when the combination of mAbs is used. The antibody concentration is the total initial concentration of antibody in the system; "mAbs" refers to tocilizumab and 10H2 together in a 1:1 concentration ratio. Simulations were performed for 24 hours after antibody dosing. The panels for [Ab] = 10 nM were presented in the main text [**Figure 7C**] and are repeated here for comparison to the other concentrations.
